# Hyoid bone position and upper airway patency: A computational finite element modeling study

**DOI:** 10.1101/2024.08.09.607294

**Authors:** Diane Salman, Jason Amatoury

**Affiliations:** Sleep and Upper Airway Research Group (SUARG), Biomedical Engineering Program, Maroun Semaan Faculty of Engineering and Architecture (MSFEA), American University of Beirut, Beirut, Lebanon

**Keywords:** finite element analysis, upper airway surgery, obstructive sleep apnea, OSA, simulation, tissue mechanics, pharynx

## Abstract

**Background and Objectives:** The hyoid bone’s inferior baseline position in obstructive sleep apnea (OSA) has led to surgical hyoid repositioning treatment, yet outcomes vary widely. The influence of baseline hyoid position (phenotype) and surgical hyoid repositioning on upper airway function remains unclear. We aimed to investigate their impact on the upper airway using computational modeling.

**Methods:** A validated finite element model of the rabbit upper airway was advanced and used to simulate changes in baseline hyoid position and surgical hyoid repositioning, alone and in combination. The hyoid was displaced in cranial, caudal, anterior, anterior-cranial and anterior-caudal directions from 1-4mm. Model outcomes included upper airway collapsibility, measured using closing pressure (Pclose), cross-sectional area (CSA) and soft tissue mechanics (stress and strain).

**Results:** Graded baseline hyoid position increments increased Pclose for all directions, and up to 29-43% at 4mm (relative to the original baseline hyoid position). Anterior-based surgical hyoid repositioning decreased Pclose (∼-115% at 4mm) and increased ΔCSA (∼+35% at 4mm). Cranial surgical hyoid repositioning decreased ΔPclose (−29%), minimally affecting CSA. Caudal surgical hyoid repositioning increased ΔPclose (+27%) and decreased ΔCSA (−7%). Anterior-cranial and anterior-caudal surgical hyoid repositioning produced the highest stresses and strains. Surgical hyoid repositioning effects on upper airway outcomes were dependent on baseline hyoid position, with more caudal baseline hyoid positions leading to less effective surgeries.

**Conclusions:** Baseline hyoid position (phenotype) and surgical hyoid repositioning both alter upper airway outcomes, with effects dependent on hyoid displacement direction and magnitude. Baseline hyoid position influences the effectiveness of surgical hyoid repositioning in reducing upper airway collapsibility. These findings provide further insights into the hyoid’s role in upper airway patency and suggest that considering the hyoid’s baseline position and surgical repositioning direction/increment may help improve hyoid surgeries for OSA treatment.

## INTRODUCTION

The hyoid bone, a mobile and central structure within the upper airway, acts as an anchor to several upper airway dilator muscles that play an important role in keeping the upper airway open during sleep. Repeated narrowing or complete collapse of the airway during sleep is characteristic of obstructive sleep apnea (OSA), a highly prevalent disorder associated with serious health consequences such as cardiovascular disease and neurocognitive impairment (1–3). A notable anatomical distinction between individuals with OSA and those without is a more inferior, or caudally located hyoid bone in individuals with OSA (4). A lower hyoid position is thought to contribute to increased upper airway collapsibility and greater OSA severity (5, 6). Efforts to treat OSA have involved surgical hyoid repositioning therapies, aiming to advance the hyoid bone and suspend it either to the mandible (hyomandibular suspension) or thyroid cartilage (hyothyroidopexy) (4, 5). While these surgical procedures have resulted in a significant decrease in the apnea-hypopnea index (AHI) in OSA, their outcomes are unpredictable and highly variable with success rates ranging from 17% to 78% (6, 7). This is likely due, at least in part, to the fact that the baseline hyoid position (phenotype) is not taken into consideration in these procedures and that the precise direction and magnitude of repositioning is not individually prescribed to achieve optimal outcomes.

Anterior hyoid advancement has been shown to considerably reduce upper airway resistance to airflow in animals (8, 9) as well as human cadavers (10). Moreover, using an anesthetized rabbit model, we recently quantified the effect of different surgical hyoid repositioning displacement magnitudes and directions on upper airway collapsibility (11). The study demonstrated that the influence of surgical hyoid repositioning on upper airway collapsibility is both direction and magnitude dependent and that anterior-based repositioning directions have the greatest effect on reducing collapsibility. However, the influence of surgical hyoid bone repositioning on upper airway geometry and tissue mechanics has not been investigated. Additionally, and importantly, the precise influence of baseline hyoid baseline (or phenotype) alone on upper airway collapse remains unknown. Finally, the impact of baseline hyoid position on the effectiveness of surgical hyoid repositioning has also not been studied.

To directly investigate the influence of baseline hyoid position on upper airway patency is not possible using physiological models. Displacing the hyoid in these models can only be accomplished by surgical repositioning and subsequent fixation, which would alter tissue mechanical properties and indeed limit hyoid movement. Accordingly, a computational model is required for this investigation.

Computational finite element modeling is particularly useful in the case of the upper airway given its complex anatomy. Indeed, the finite element method has been extensively employed for biomedical applications (12), including the upper airway (13–16). We have previously developed and validated a two-dimensional (2D) computational finite element model of the passive rabbit upper airway (17), which includes the hyoid bone, major upper airway soft and bony tissues, as well as the passive influence of many upper airway muscles. The model was developed alongside a physiological anaesthetized rabbit model (18, 19), which allows for continuous model enhancement and the ability to directly validate model predictions. Indeed, the use of the rabbit as a model for the upper airway is ideal given anatomical similarities between the rabbit and human upper airway, particularly the mobility of the hyoid bone (18, 20). Furthermore, rabbit upper airway outcomes have repeatedly demonstrated similarities to the human circumstance (9, 11, 21–31). This computational model is uniquely capable of predicting upper airway geometry and soft tissue mechanical changes in response to mandibular advancement and tracheal displacement. Notably, no other upper airway model, to our knowledge, boasts a comparable level of development, validation, and functionality, including the ability to manipulate the hyoid bone. However, the effect of altering hyoid position was not considered and the model lacked the ability to simulate negative intraluminal pressures to study upper airway collapsibility.

The aim of the current study is to advance our previously developed computational model of the upper airway (17) and investigate the influence of baseline hyoid position and surgical hyoid repositioning, independently and combined, on upper airway patency, collapsibility, and tissue mechanics.

## MATERIALS AND METHODS

### Overall Study Design

The previously validated 2D computational finite element model of the passive rabbit upper airway by Amatoury et al. (17) was advanced to allow hyoid position changes (baseline and surgical) and upper airway collapse. Using ANSYS Workbench (Release 19.2; Academic Research, Canonsburg, PA), the advanced model simulated 1) changes in baseline hyoid position, 2) surgical hyoid repositioning and 3) combined changes in baseline hyoid position and surgical repositioning, each with subsequent application of negative intraluminal pressure for upper airway collapsibility determination. Model outcomes included upper airway closing pressure (Pclose), upper airway cross-sectional area (CSA) and tissue mechanics (displacement, stress and strain).

### Finite Element Model Definitions

#### Model geometry, material properties, contact and boundary conditions

The finite element model geometry, material properties and boundary conditions have been described in detail previously (17), and only slight modifications were made in the current model to allow for current simulations (described below). Briefly, the model consisted of a 2D mid-sagittal representation of the rabbit upper airway, including all major upper airway structures such as the tongue, soft and hard palate, constrictor muscles, thyroid cartilage, epiglottis, mandible and hyoid bone. The rest of the soft tissues, mainly including adipose and glandular tissue, are lumped into a single structure named the tissue mass. A rectangular base plate was also added to the lower edge of the thyroid cartilage to provide a surface for the attachment of select muscles (and application of tracheal displacement in the previous study, (17)) (Figure 1A B).

**Figure 1:**
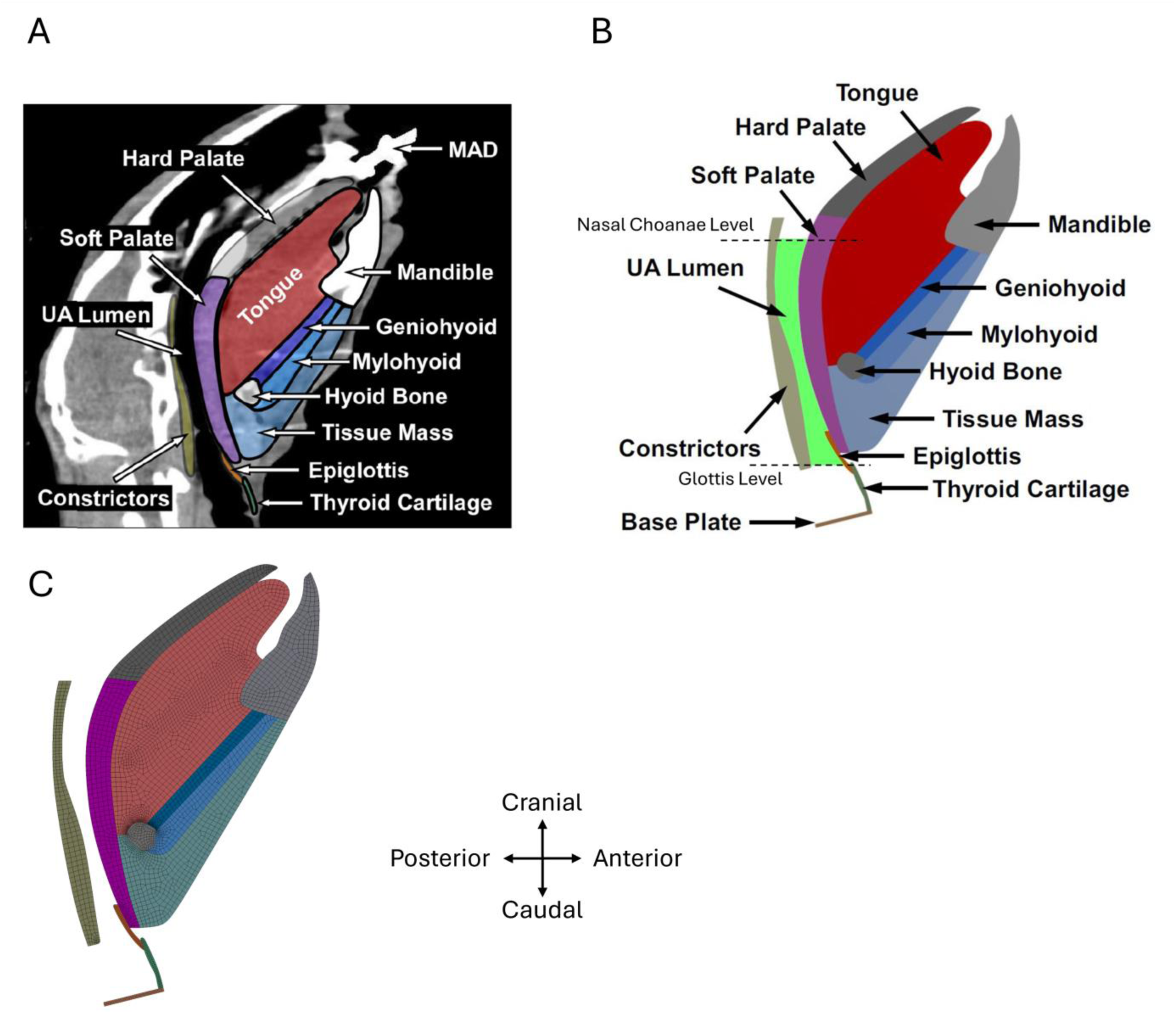
Model geometry and final mesh. (A) Midsagittal computed tomographic (CT) image with labelled and colored tissues used for FE model geometry reconstruction [adapted from Amatoury et al 2016 (17)]. (B) 2D mid-sagittal rabbit upper airway FE Model geometry including bony, cartilaginous and soft tissue structures (labelled) and upper airway lumen [highlighted in green, defined from the level of the nasal choanae to the glottis; adapted and modified from Amatoury et al 2016 (17)]. (C) FE model mesh of primarily two-dimensional 4-node quadrilateral plane strain elements. The final mesh consists of 3970 elements with 3907 quadrilateral elements and 63 triangular elements.

Bony and cartilaginous tissues were assigned incompressible linear elastic material properties (Table 1) (17). A nonlinear hyperelastic material model (Yeoh second-order strain energy function) was used for soft tissues (Table 2) (17). The passive action of pharyngeal muscles was represented using linear elastic spring connections translated into the mid-sagittal plane (Figure 2). The contact and boundary conditions were defined from anatomy and physiological experimentation (18, 19), and are summarized in Figures 3 and 4. Further details regarding the boundary conditions, contact conditions and material properties can be found in Amatoury et al. (17).

**Figure 2:**
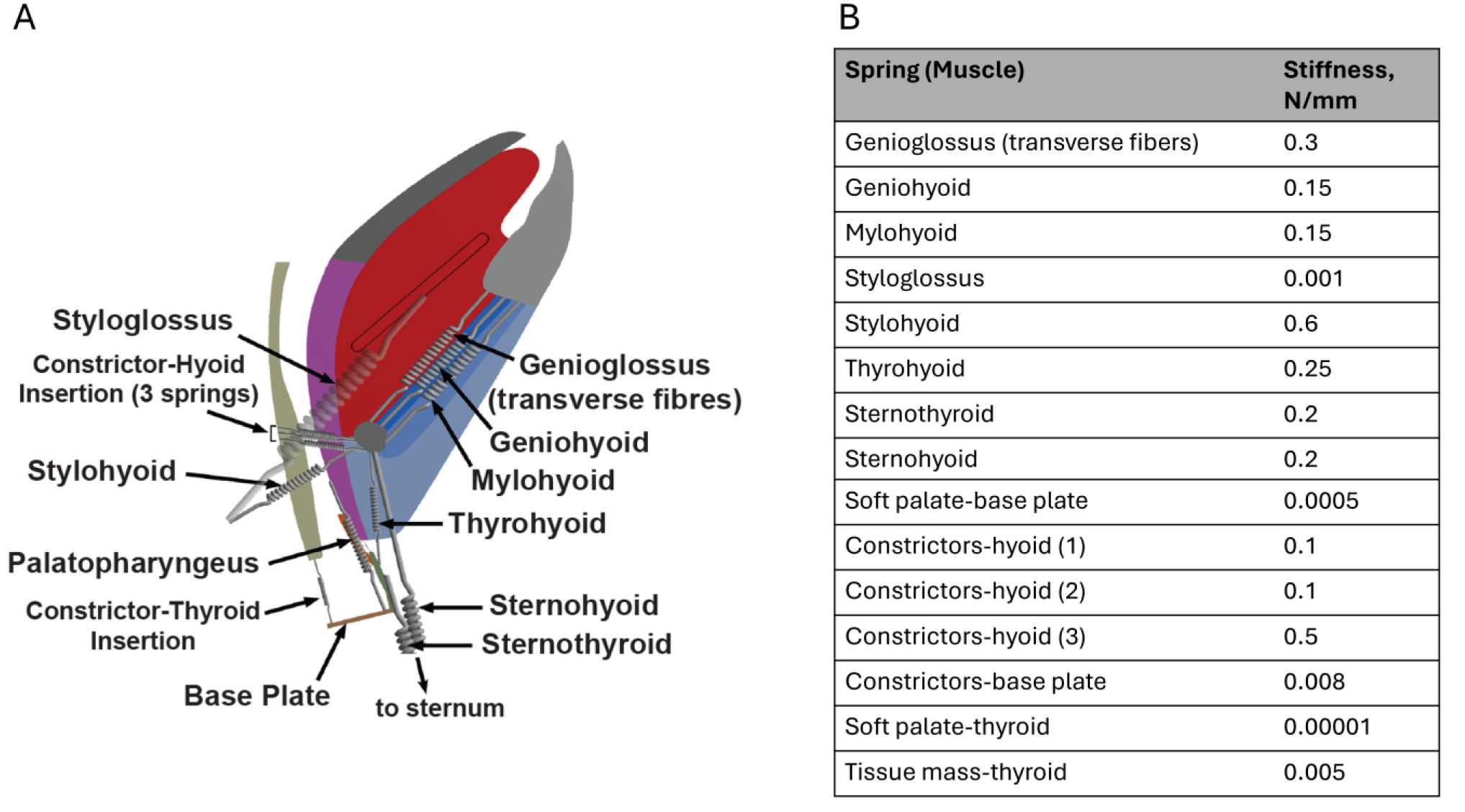
Passive pharyngeal muscle connections. The passive action of multiple pharyngeal muscles were modeled as linear elastic springs (A), with varying stiffness properties (B). Adapted and modified from Amatoury et al 2016 (17).

**Figure 3:**
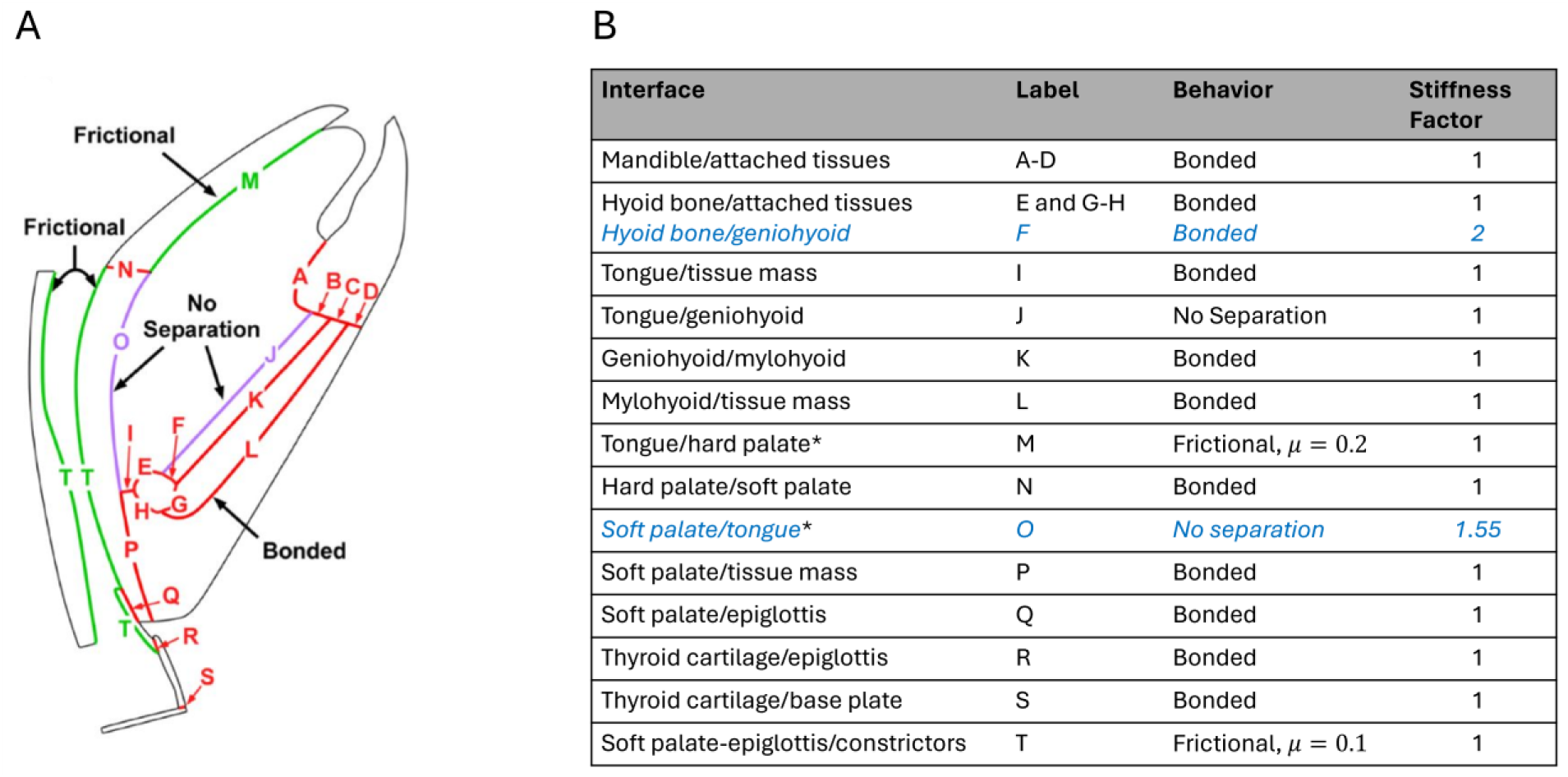
Model contact conditions. (A) Contact definitions on model geometry edges are shown with labels and colors. Red outlines represent bonded contact definitions, purple outlines no separation contacts and green the frictional contact definitions. (B) Table summarizing model contact interfaces and corresponding properties including contact behavior and normal stiffness factor (default stiffness factor is 1). Contacts in italic (and blue) represent the contacts for which the stiffness factor was modified from the original model. * indicates that the contact interface was modified from the original model. Label (second column) point to the location of interface shown in (A) Adapted and modified from Amatoury et al 2016 (17).

**Figure 4:**
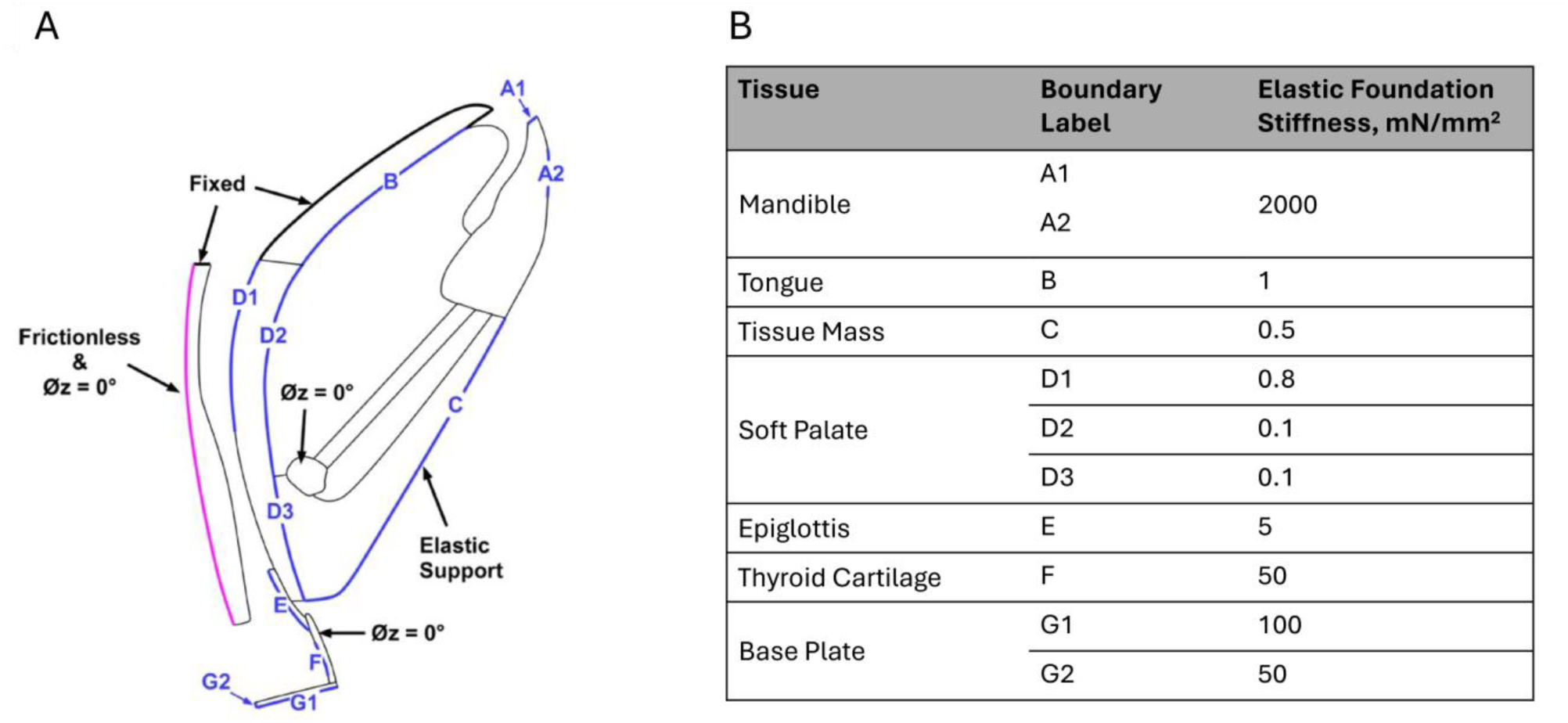
Model boundary conditions. (A) Boundary labels illustrated on the model where blue outlines indicate elastic supports (labels A1 to G1), dark black outlines indicate fixed support, pink outline indicates frictionless support and no rotation condition (Øz = 0°) is applied to the constrictors, thyroid cartilage and hyoid bone. (B) Corresponding elastic constraints properties (stiffness) for the different boundaries, with the boundary label pointing to the location in (A). Adapted and modified from Amatoury et al 2016 (17).

**Table 1:**
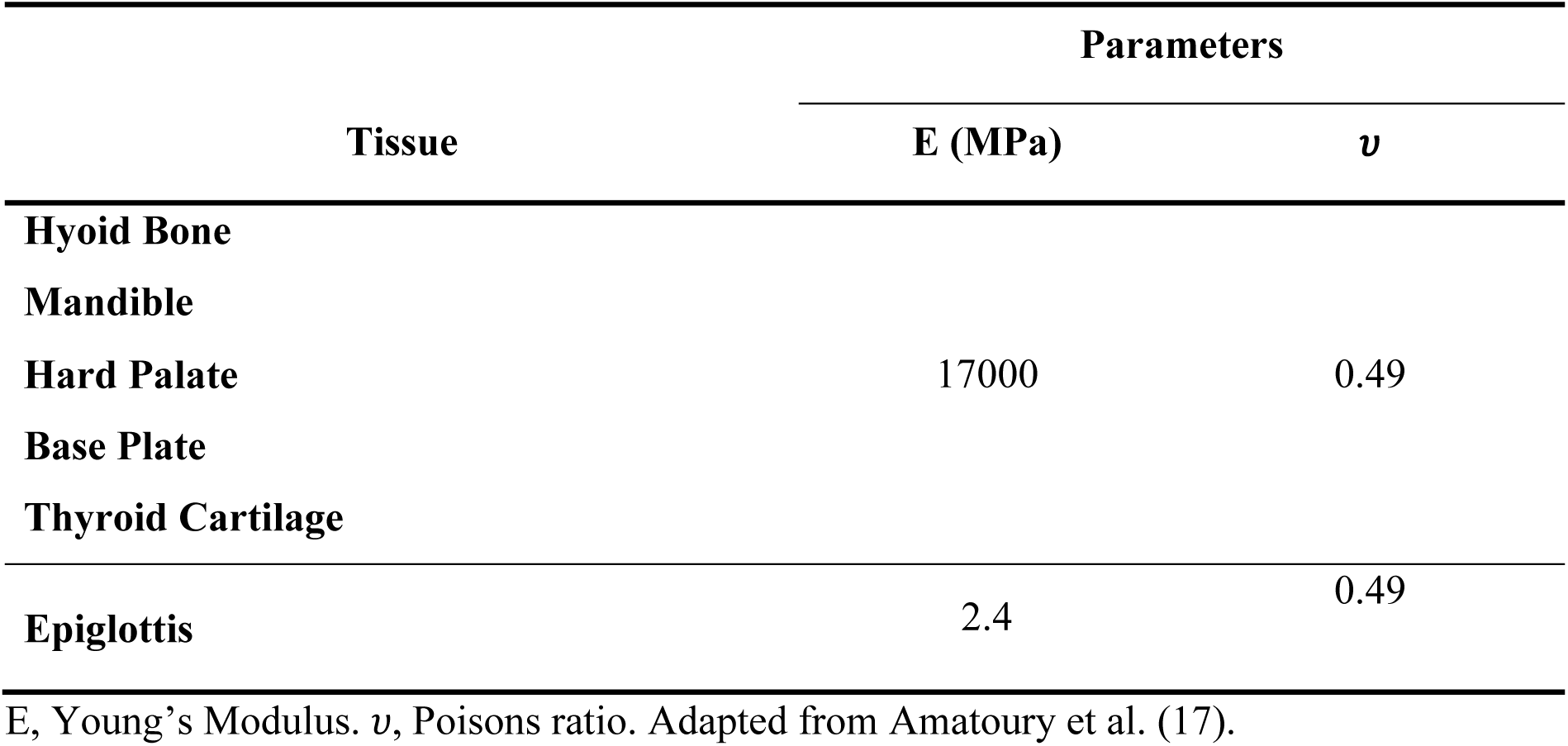
Linear elastic material for bony and cartilaginous tissues.

**Table 2:**
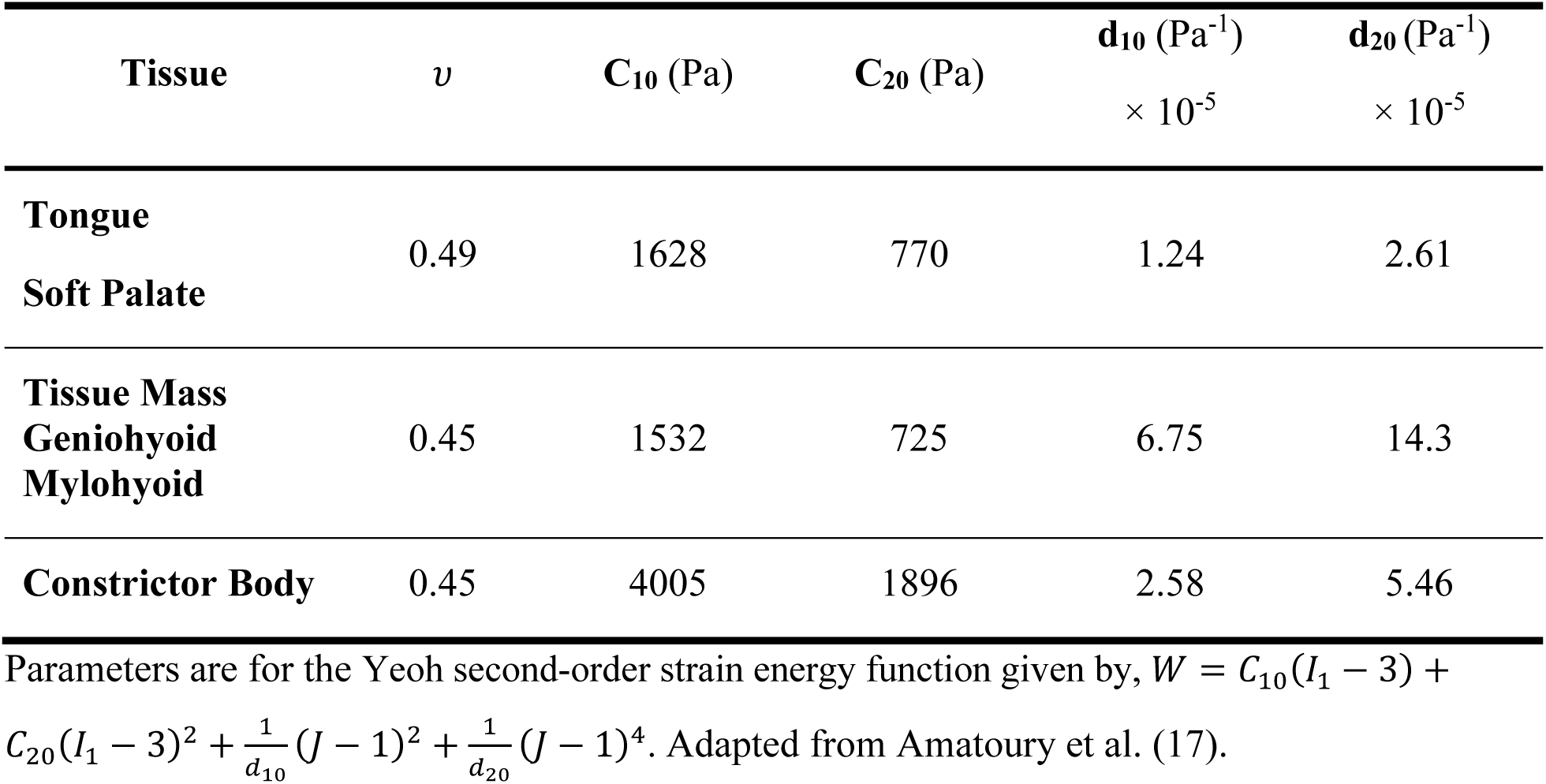
Nonlinear hyperelastic model parameters (Yeoh second-order) for soft tissues.

### Model redevelopment and advancement

Slight geometrical modifications were made to the original model to allow for the geometric representation of different baseline hyoid positions (phenotypes) in the current model. This primarily involved linking the hyoid geometrically to the tissues surrounding it such that a baseline shift in hyoid position would also cause all attached structures to shift with it. For example, an anterior shift in the hyoid baseline position also leads to a forward extension of the base of the tongue and a change in the angle and length of the geniohyoid and mylohyoid surfaces, in addition to movement of hyoid attached springs.

Simulating upper airway collapse required the adjustment of select boundary conditions, mesh and analysis settings from the original model. The no-separation contact originally defined between the soft palate and tongue (Labeled O in Figure 3) and the frictional contact between the hard palate and tongue (Labeled M in Figure 3) were redefined to include the whole upper boundary of the tongue to adapt to the large tongue displacements during Pua simulations. Based on the Newton-Raphson residuals, adjustments to the stiffness factor of some contacts were also made.

### Mesh

The mesh was generated using ANSYS meshing capabilities. Two-dimensional 4-node quadrilateral plane strain elements were primarily used, which adapt well to large deformation produced by the hyperelastic and nearly incompressible soft tissues represented in the current model. The default pure displacement element formulation was applied and the element technology used was the simplified enhanced strain formulation to prevent shear locking. Moreover, the mesh was refined at the edges around the hyoid bone since higher strains were expected in this area. Additional mesh refinement was performed at the edge between the hyoid and the geniohyoid surface and the edge between the hyoid and tongue. The final mesh consisted of 3970 elements with 3907 quadrilateral elements and 63 triangular elements (Figure 1C; see Appendix A for mesh convergence).

### Hyoid geometry changes, loading conditions and simulation protocol

Baseline hyoid position changes were achieved by altering the hyoid bone’s geometrical position, i.e. in the model geometry. Connected hyoid tissues/springs would move with the hyoid, but mechanical properties of these tissue/spring connections (e.g. stiffness) would not change. Surgical hyoid repositioning was modeled by applying a displacement load at the centroid of the hyoid bone surface, which was then fixed in the new position. In this case, tissue connections would be stretched or compressed, depending on the direction of displacement. In both baseline and surgical cases, the hyoid was moved within the midsagittal plane along cranial, caudal, anterior, anterior-caudal (45°) and anterior-cranial (45°) directions (Figure 5A). For each direction, the hyoid was repositioned (baseline geometry change or displacement load) from 1-4mm in 1mm increments.

**Figure 5:**
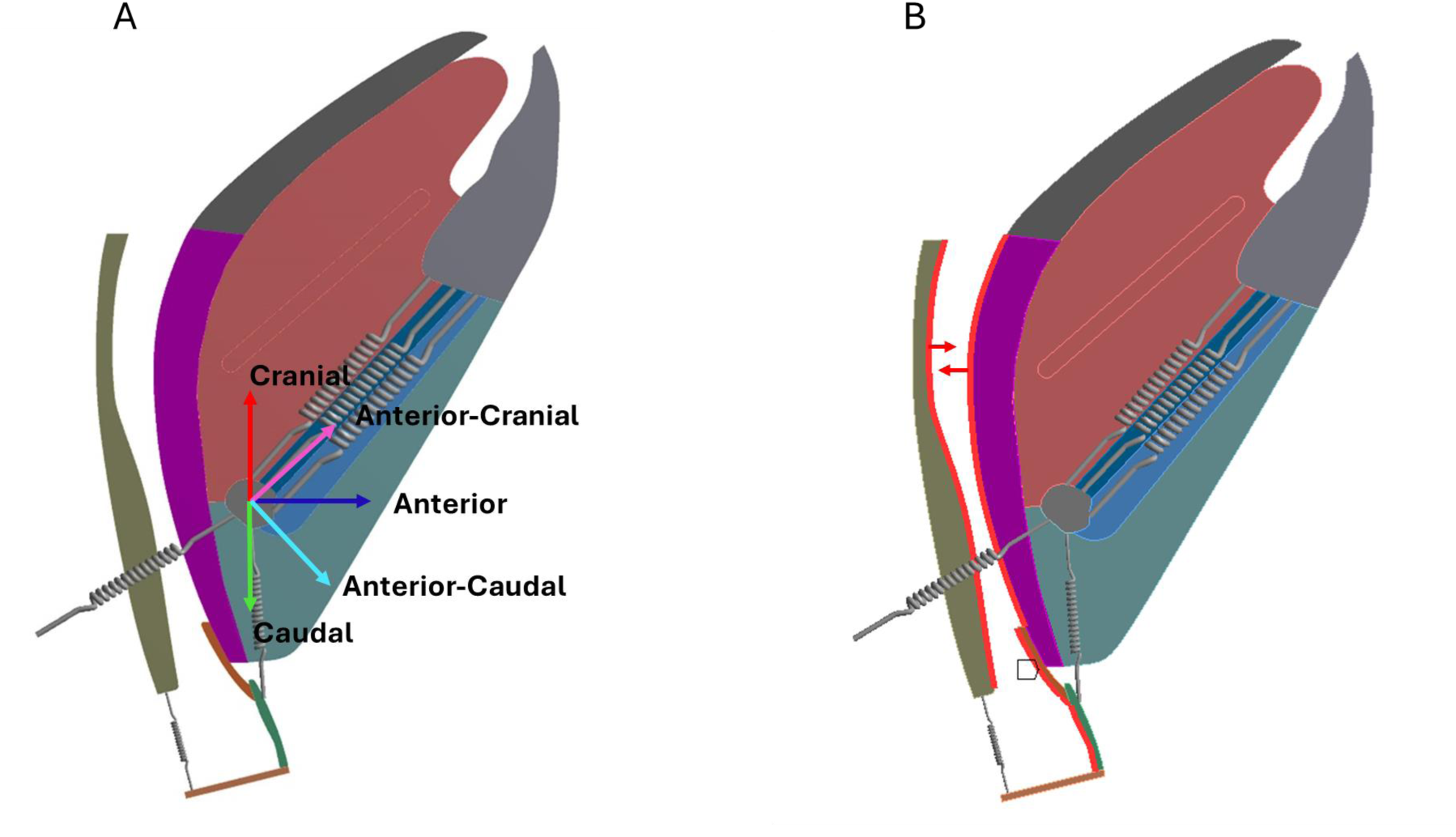
Hyoid displacement and negative intraluminal pressure load. (A) Directions of hyoid displacement for baseline hyoid position changes and surgical hyoid repositioning simulations. (B) Negative intraluminal pressure load was applied as a uniformly distributed negative pressure on opposite airway walls (constrictors and soft palate/epiglottis/thyroid).

Negative intraluminal pressure (Pua) was applied by a uniformly distributed negative pressure load across opposite walls of the upper airway, delineated by the constrictors posteriorly and the soft palate, epiglottis and thyroid anteriorly (Figure 5B). This load was simulated to determine upper airway collapsibility (described further below). In each simulation, a Pua load magnitude of −9000 Pa (−91.8 cmH_2_O) was applied, a pressure selected to ensure collapse across all simulation conditions, facilitating precise pressure determination with the substeps utilized in the finite element analysis (see below). The absolute Pua is greater than that which would be applied experimentally, but we are concerned with the change in pressure within each intervention rather than the absolute value. This limitation with the absolute Pua load is addressed in discussion.

The simulation protocol involved the following: 1) for each baseline hyoid position or 2) surgical hyoid repositioning load, a Pua load was applied. Furthermore, to investigate the impact of hyoid baseline position changes on the effectiveness of surgical hyoid repositioning in decreasing upper airway collapsibility, 3) surgical hyoid repositioning was applied at altered baseline hyoid positions, followed by a Pua load. The surgical hyoid displacements simulated were those that are clinically relevant, including anterior-cranial surgical hyoid repositioning, akin to hyomandibular suspension surgery, and anterior-caudal surgical repositioning, akin to hyothyroidopexy. These were performed at 0,2 and 4mm surgical repositioning, for 0, 2 and 4mm baseline hyoid position changes in all directions, including the OSA phenotypic hyoid characteristic of a more caudally located hyoid bone (32, 33). Figure 6 summarizes the simulation protocol and model outputs which are described in further detail below.

**Figure 6:**
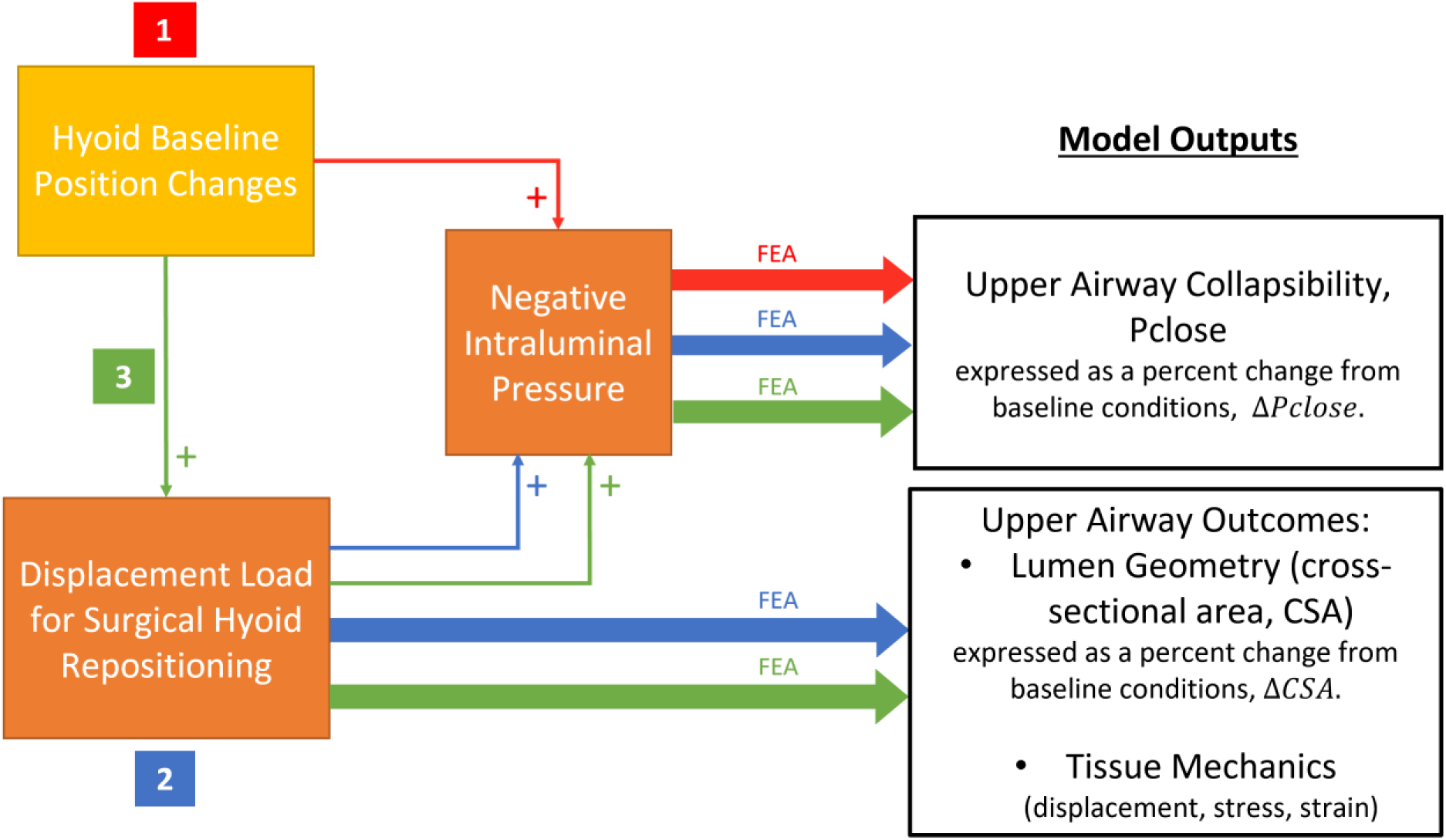
Simulation protocol and model outputs. The finite element analysis (FEA) simulations performed in the study included: 1) Hyoid baseline position changes, 2) surgical hyoid repositioning from the original baseline position, 3) surgical hyoid repositioning starting from different hyoid baseline positions. Negative intraluminal pressure was applied following each 1-3. Outcomes quantified include upper closing pressure (Pclose), airway lumen cross-sectional area (CSA) and tissue mechanics.

### Finite element analysis settings

The finite element analysis was performed under a static structural system with 2D plane strain conditions using the ANSYS Mechanical solver. Large deformation effects were enabled due to the presence of non-linear hyperelastic materials and expected high strains. The hyoid displacement load (for surgical hyoid repositioning) was applied over a single step with program controlled substeps. The Pua load was also applied over a single load step, but with 80 substeps for the purpose of quantifying Pclose. That is, Pclose is dependent on the number of substeps (or increments in Pua), where the Pclose measurement error is defined as half the smallest Pua increment. After testing for different number of substeps, 80 was found to be the ideal number for solution convergence as well as relative accuracy of Pclose determination with a measurement uncertainty of ∼±1.4% relative to Pclose at baseline in the current model (see below for Pclose quantification).

### Model Outputs

#### Upper airway collapsibility (closing pressure, Pclose)

The collapsibility of the upper airway was quantified using Pclose, the pressure at which the upper airway closes. Pclose was taken as the pressure at which the opposing upper airway walls first come into direct contact. This was achieved by monitoring the contact between the anterior side of the constrictors and the posterior boundaries of the soft palate, epiglottis and thyroid cartilage throughout the Pua load step using the post-processing contact tool in ANSYS. The substep at which the first non-zero penetration value occurs (i.e. walls in contact) was taken as the point of collapse. Pclose results were represented as the percent change from original baseline (hyoid position = 0mm; ΔPclose). For combined changes in baseline hyoid position and surgical repositioning changes simulations, Pclose results were represented as the precent change from the respective baseline hyoid position (surgical repositioning = 0mm; Δ’Pclose).

### Upper airway lumen cross sectional area (CSA)

The changes in upper airway lumen cross-sectional area (CSA) in response to different hyoid displacement loads were assessed by exporting the deformed model geometry into SOLIDWORKS (Version 2016, Dassault Systèmes). The upper airway contour was created from the level of the nasal choanae to the glottis (Figure 1B) and its CSA determined, as previously described (17–19). CSA outcomes were presented as a percent change from the baseline undeformed airway (hyoid position=0mm; ΔCSA). CSA was computed for select increments of surgical hyoid repositioning, 2 and 4mm, comparable to the hyoid displacement magnitudes applied clinically in surgical hyoid repositioning procedures (32, 33).

### Tissue displacement, stress and strain

The displacements of upper airway structures in response to the applied loads were displayed as color-coded displacement contours on the deformed mesh. The equivalent (von Mises) stress and strain distributions in the upper airway soft tissues were also visualized as color-coded contour maps. Similar to CSA, the resultant displacement, stress and strain distributions of upper airway tissues were computed for select increments of surgical hyoid repositioning (2 and 4mm)

### Model Verification

To verify that the model was independent of the mesh density, a mesh convergence study was performed using the output of tissue displacement at the site of airway collapse. Additional details can be found in Appendix A.

### Model Validation

The model was validated for Pclose outcomes with surgical hyoid repositioning against experimental data obtained by Samaha et al., who performed surgical hyoid repositioning using our complementary anaesthetized rabbit model (11). The experimental data were originally presented as an absolute change from baseline (11), but were recalculated as percent change in order to compare with the outcomes of the current study. Specifically, for each direction (cranial, caudal, anterior, anterior-cranial, anterior-caudal) and increment (0, 1, 2, 3 and 4mm) of hyoid displacement, the percent change in Pclose with respect to the baseline Pclose (ΔPclose, %) obtained from model simulations were compared to the mean ± 95% confidence interval of these outcomes obtained experimentally. In total, the model was compared against 20 experimental outcomes. The finite element model was considered in strong agreement with experiments if Pclose outputs fell within the 95% confidence limits for the relevant experimental data.

## RESULTS

### 1. Baseline hyoid position

All directions and increments of baseline hyoid position changes lead to an increase in Pclose, indicating greater collapsibility, as seen in Figure 7. The gradual increase in ΔPclose for all directions reached up to 29-43% at 4mm, with the greatest change occurring for anterior-cranial and anterior-caudal direction. The caudal direction closely followed, particularly at lower increments (Figure 7).

**Figure 7:**
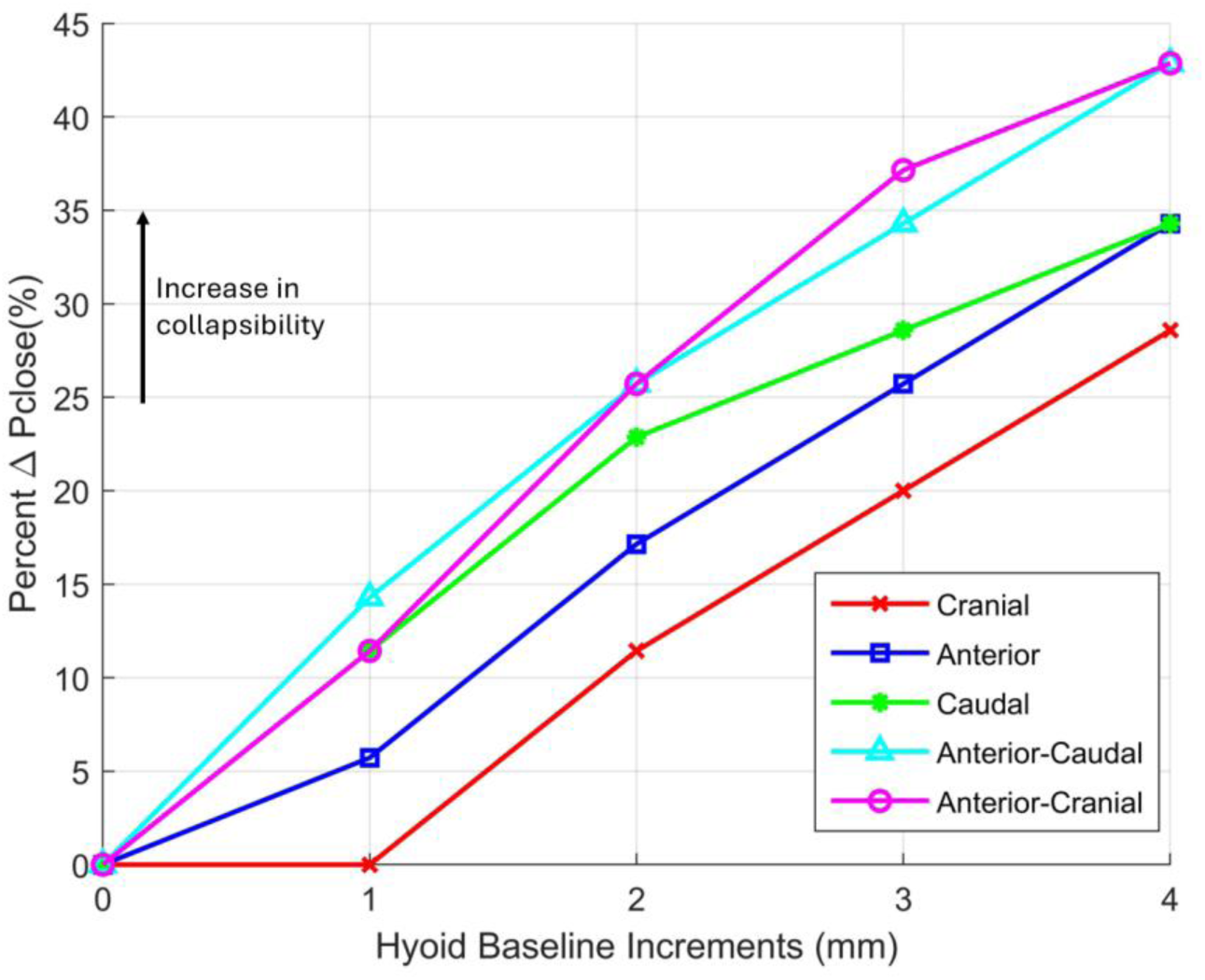
Upper airway collapsibility vs baseline hyoid position. Plot showing the percent change in closing pressure (ΔPclose) when the baseline hyoid position is incremented in the cranial, anterior, caudal, anterior-cranial and anterior-caudal directions (different symbols and colors). Incremental baseline hyoid position changes resulted in an increase in ΔPclose for all directions, indicating increased upper airway collapsibility.

### 2. Surgical hyoid repositioning

#### Pclose

The change in Pclose with surgical hyoid repositioning from the original baseline hyoid position were dependent on both the direction and magnitude of surgical hyoid repositioning, as shown in Figure 8. The largest decrease in Pclose was achieved with anterior-based surgical repositioning directions, ranging between approximately −110 to −120% at 4mm. Cranial surgical hyoid repositioning also resulted in a relatively slight decrease in Pclose of up to −29% at 4mm. However, caudal surgical hyoid repositioning simulations had little to no effect on Pclose at 1 and 2mm increments, then subsequently caused a slight increase in Pclose of up to 27% at 4mm.

**Figure 8:**
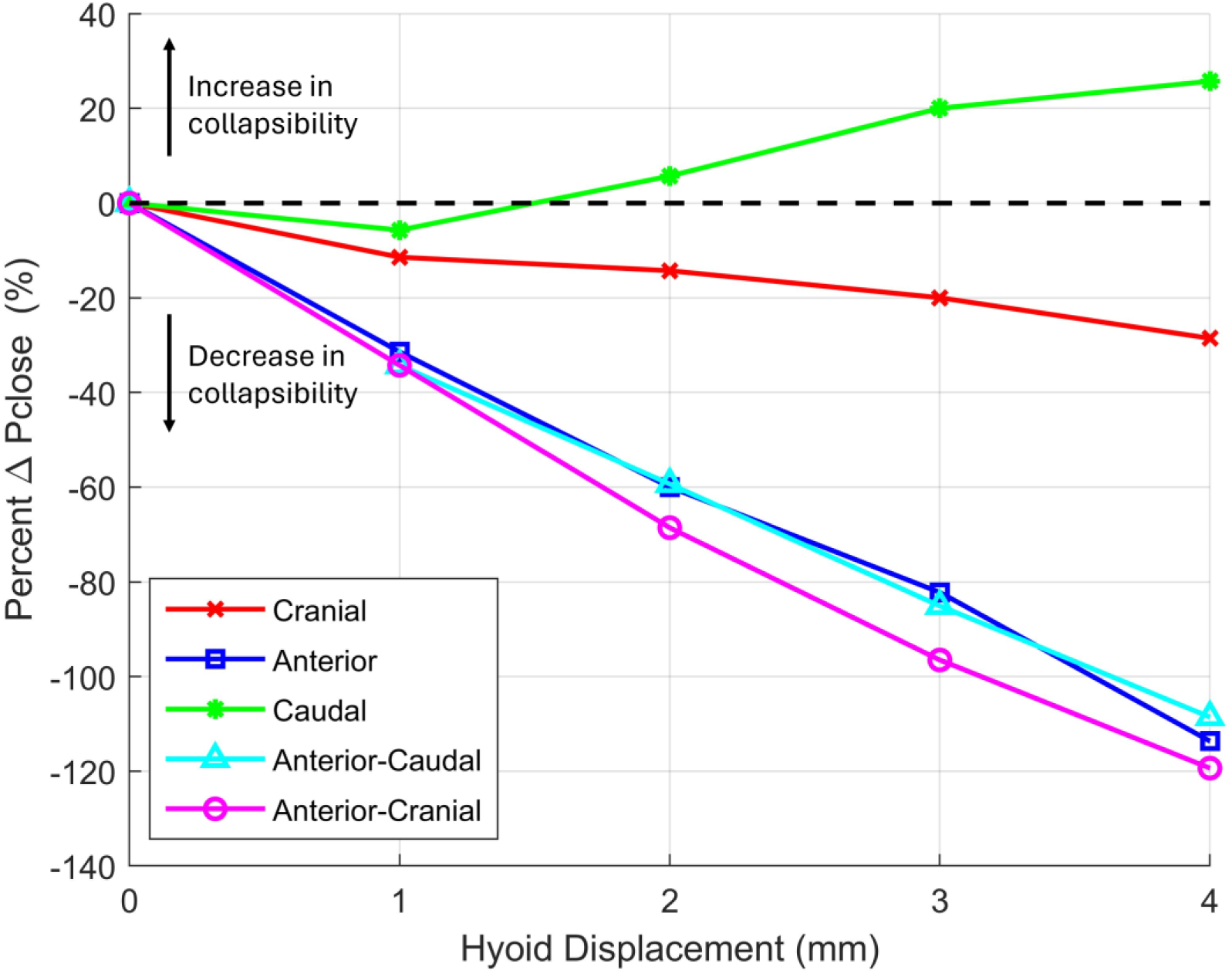
Upper airway collapsibility vs surgical hyoid repositioning. Plot representing percent change in closing pressure (ΔPclose) with increasing surgical hyoid repositioning from the original hyoid baseline position in the cranial, anterior, caudal, anterior-caudal and anterior-cranial directions (different symbols and colors). The change in Pclose is dependent on both the direction and magnitude of hyoid displacement. Anterior-based directions led to the largest decrease in ΔPclose.

#### Model validation for Pclose

Pclose outcomes were successfully validated against experimental data from the animal study by Samaha et. al. (11) (see Figure 9). FE model Pclose results were within the 95% confidence interval of the experimental Pclose outcomes (strong agreement) for all directions and increments of surgical hyoid repositioning, except caudal repositioning at 1 and 4mm, which were just outside the 95% confidence interval (Figure 9). Thus, the computational model can besaid to successfully predict changes in upper airway collapsibility produced experimentally by surgical hyoid repositioning procedures.

**Figure 9:**
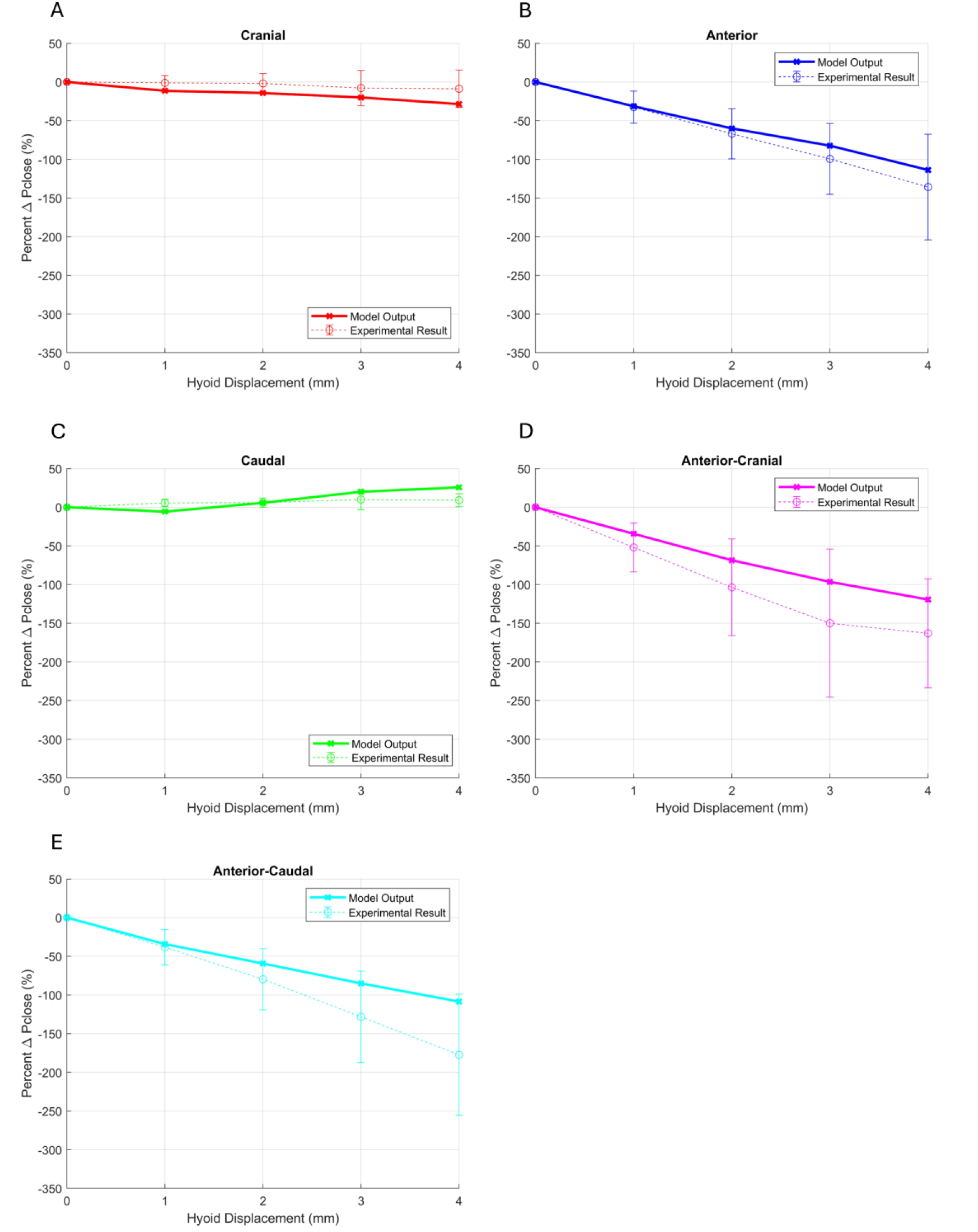
Validation of upper airway collapsibility model predictions. The percent change in closing pressure (ΔPclose) produced by surgical hyoid repositioning simulations vs. experimental outcomes from Samaha et al. (11). Plots show the experimental mean outcomes with 95% confidence interval bars and model output for surgical hyoid repositioning in the cranial (A), anterior (B), caudal (C), anterior-cranial (D) and anterior-caudal (E) directions. Model ΔPclose outputs were within the 95% confidence interval of experimental results except for select increments of caudal and cranial hyoid repositioning interventions.

#### Tissue displacement and CSA

Tissue displacement distributions were dependent on the repositioning direction. Changes in repositioning increments in the same direction only led to higher magnitudes of displacement while maintaining a similar distribution across tissues. Figure 10A shows the total displacement contour maps at the maximum 4mm hyoid repositioning increment in all directions. Soft tissue structures, including the tongue, soft palate, tissue mass, geniohyoid and mylohyoid, were displaced in response to hyoid repositioning with the largest displacements being around the hyoid bone. Anterior-cranial and anterior-caudal repositioning directions produced the largest tissue displacements. More specifically, anterior-based directions resulted in large anterior displacement of the soft palate region behind the hyoid bone which visibly widened the airway lumen in that area.

**Figure 10:**
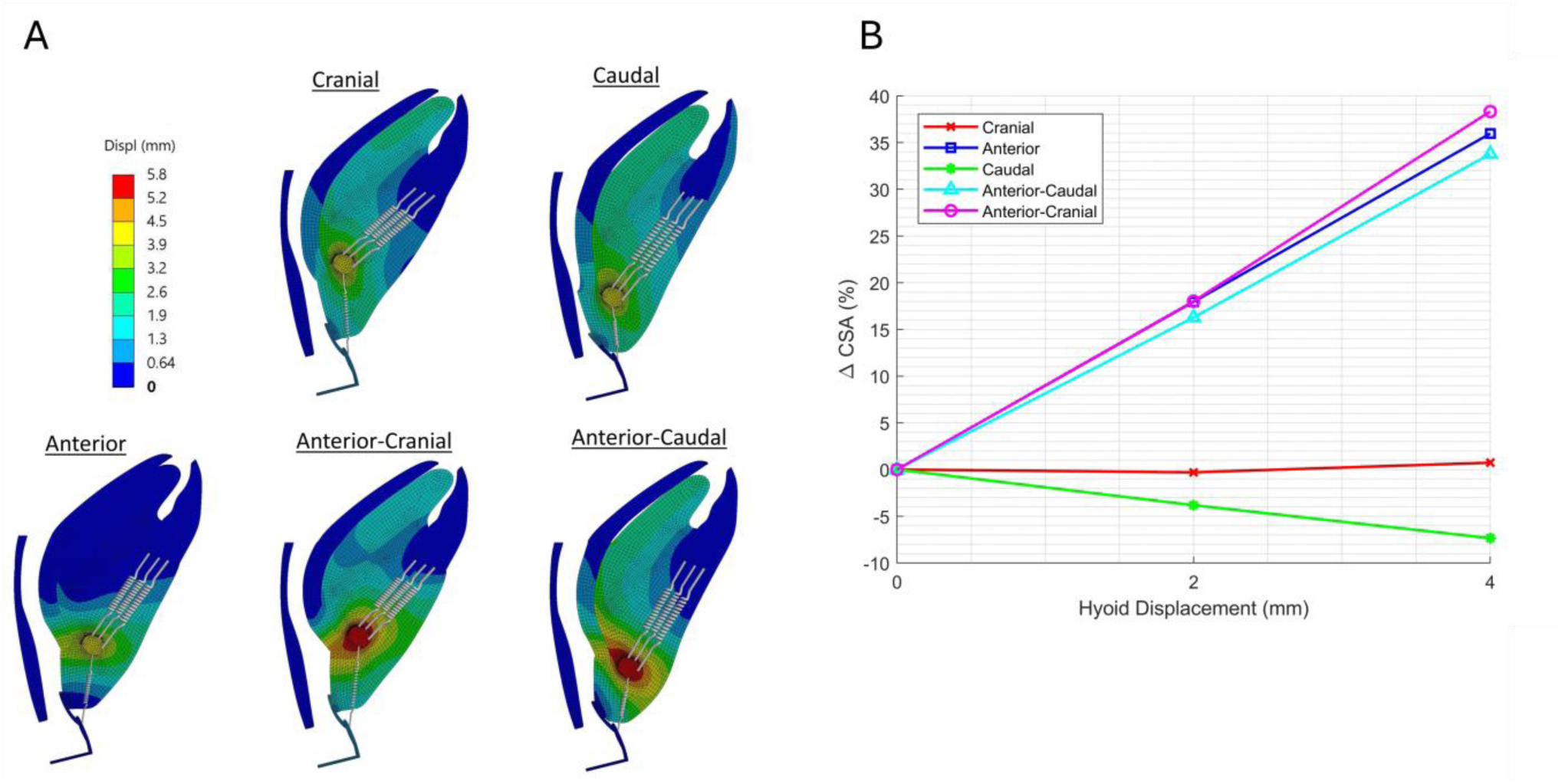
Tissue displacement and upper airway cross-sectional area (CSA) vs surgical hyoid repositioning. (A) Color-coded resultant tissue displacement (Displ) fields shown at the maximum 4mm surgical hyoid repositioning increment in cranial, caudal, anterior, anterior-cranial and anterior-caudal directions from the original baseline hyoid position. Note only select springs are shown for improved visualization. (B) Percent change in CSA (ΔCSA) from original baseline hyoid position with surgical hyoid repositioning in all directions. Hyoid repositioning in the anterior-based directions led to the greatest increase in ΔCSA, with little change cranially and a slight decrease caudally.

Anterior-based hyoid repositioning directions led to the largest increase in upper airway lumen CSA, ranging between 30 and 40% at 4mm (Figure 10B). Cranial repositioning resulted in a negligeable change in CSA whereas caudal repositioning caused a minor reduction in CSA (−7% at 4mm increment).

#### Stress and strain

Stress and strain distributions vary depending on the direction of surgical hyoid repositioning (Figure 11). Anterior-cranial repositioning generated the highest stresses and strains across soft tissue areas extending from the soft palate region inferior to the hyoid bone to the boundary with the mandible, including the tongue, geniohyoid and mylohyoid areas, and tissue mass.

**Figure 11:**
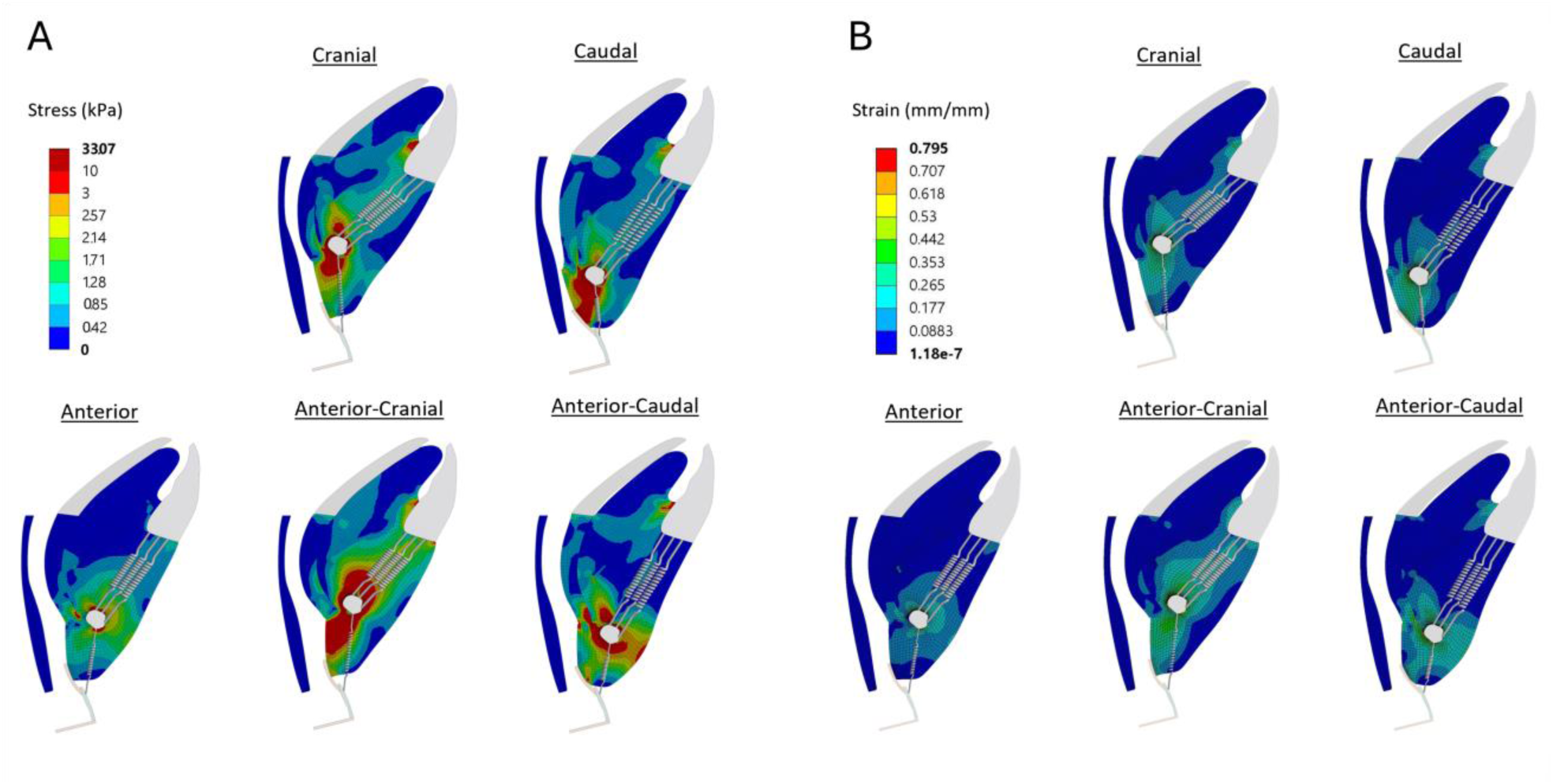
Soft tissue stress and strain vs surgical hyoid repositioning. Stress (A) and strain (B) distribution results for 4mm surgical hyoid repositioning from the original baseline hyoid position in the cranial, caudal, anterior, anterior-cranial and anterior-caudal directions. Stress and strain distributions vary depending on the direction of hyoid repositioning. Anterior-cranial repositioning generated the highest stress and strain distributions across soft tissue areas, particularly around the hyoid bone. Note only select spring are shown for improved visualization.

### 3. Combined baseline hyoid position changes and surgical hyoid repositioning in anterior-cranial and anterior-caudal directions

#### Pclose

Figure 12 shows the change in Pclose for simulations involving 2 and 4mm anterior-cranial and anterior-caudal surgical hyoid repositioning from different baseline hyoid positions. The simulations revealed that alterations in Pclose due to surgical hyoid repositioning are dependent on the initial baseline hyoid position. Specifically, when the hyoid was in its original baseline position, 2mm anterior-cranial surgical repositioning led to a decrease in Pclose by approximately −70% compared to baseline. For a hyoid that is positioned in more anterior, cranial or anterior-cranial directions at baseline, Pclose was improved to between −80 to −120% (depending on direction/increment) with 2mm anterior-cranial surgical repositioning. However, for clinically relevant caudal baseline position of 2 or 4mm (OSA phenotype), anterior-cranial surgical repositioning resulted in only a 43% decrease in Pclose at 4mm. Similarly, a 2mm anterior-caudal surgical repositioning resulted in a ∼60% decrease in Pclose at the original baseline position compared to a 52% decrease at a 4mm caudal baseline position. In essence, simulations of 2mm anterior-cranial and anterior-caudal surgical hyoid repositioning were approximately 25% and 8% less effective, respectively, in decreasing Pclose for a 4mm caudal baseline hyoid position.

**Figure 12:**
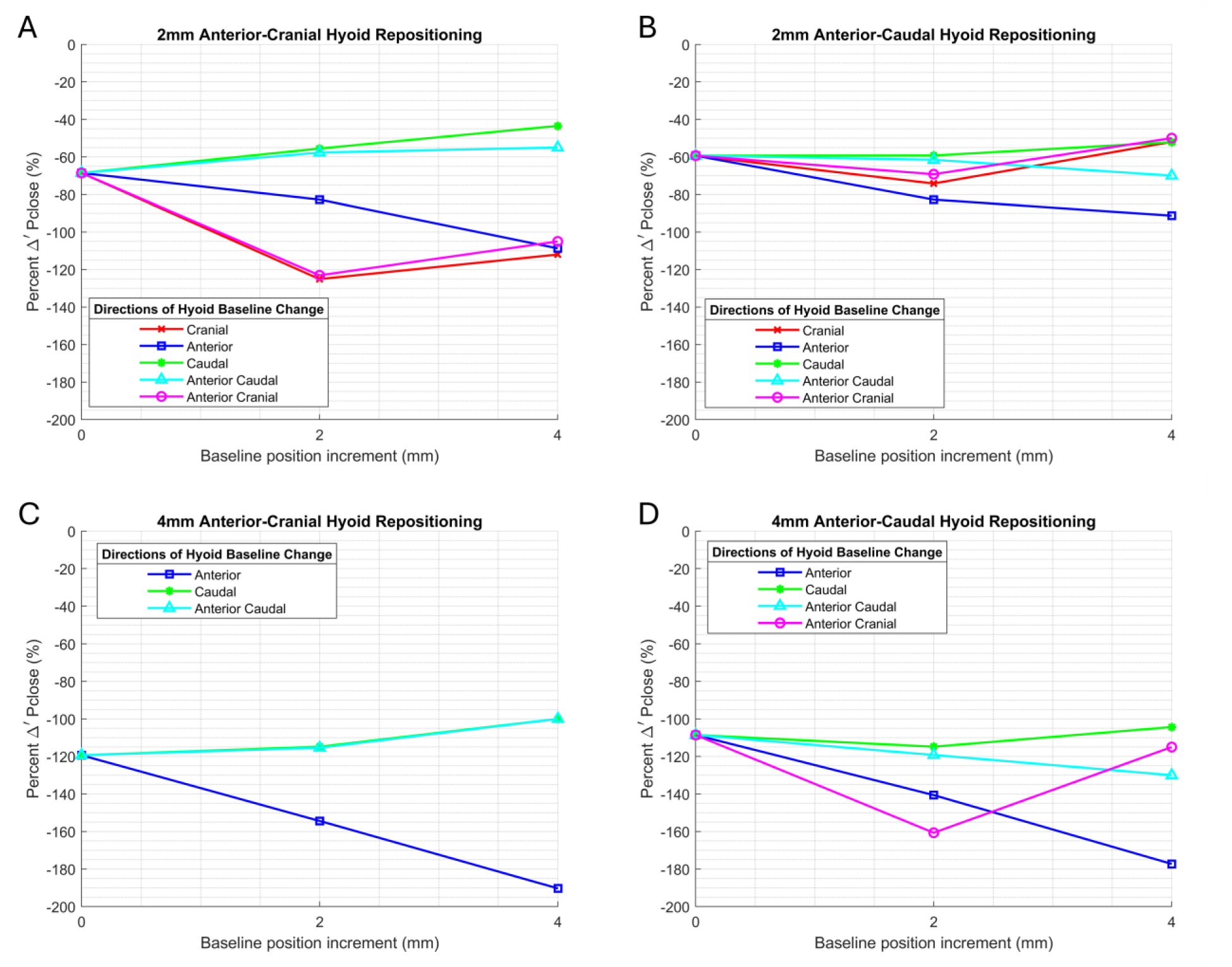
Surgical hyoid repositioning from different baseline hyoid positions vs. upper airway closing pressure (Pclose). Influence of (A) 2mm and (C) 4mm anterior-cranial and (B) 2mm and (D) 4mm anterior-caudal surgical hyoid repositioning for 0, 2 and 4mm baseline hyoid position changes in all directions on Δ′Pclose [ΔPclose relative to the respective (altered) hyoid baseline position Pclose (surgical hyoid repositioning = 0mm)]. Anterior-cranial and anterior-caudal surgical hyoid repositioning are akin to hyomandibular suspension and hyothyroidopexy surgical procedures, respectively. Surgical hyoid repositioning outcomes for Δ′Pclose are shown to be dependent on the baseline hyoid position. In particular, the clinically relevant case of a more caudal hyoid baseline position, the same surgical hyoid repositioning simulation results in a higher Δ′Pclose (less reduction in collapsibility) for a more caudally positioned hyoid (full circles; green line). Note missing data points/lines for 4mm surgical repositioning are where model convergence was not achieved.

With 4mm surgically repositioning, the model was unable to converge on a solution for Pclose in the cranial and/or anterior-cranial baseline hyoid position directions, however, the trend in the outcomes for the available 3-4 directions were similar to the 2mm repositioning case (Figure 12). For the particularly relevant caudal position, 4mm anterior-cranial and anterior-caudal surgical hyoid repositioning directions were ∼19% and 4% less effective in decreasing Pclose, respectively.

#### Tissue displacement and CSA

Tissue displacement and CSA results were examined for the clinically relevant caudal hyoid baseline positions (similar to OSA phenotype) combined with anterior-cranial and anterior-caudal surgical hyoid repositioning simulations, as shown in Figure 13. Tissue displacement distributions were dependent on the direction of surgical hyoid repositioning (Figure 13A). While anterior-cranial repositioning led to slightly larger displacements, anterior-caudal repositioning deformed a wider portion of the anterior upper airway wall. However, in both cases, largest soft tissue displacements were observed around the hyoid bone and mainly in the inferior portion of the soft palate leading to the widening of the lower section of the upper airway lumen. In fact, both anterior-cranial and anterior-caudal surgical hyoid repositioning simulations significantly increased the airway lumen CSA by approximately 35% (Figure 13B,C). Changes in the increment of baseline hyoid position produced only a small difference in ΔCSA.

**Figure 13:**
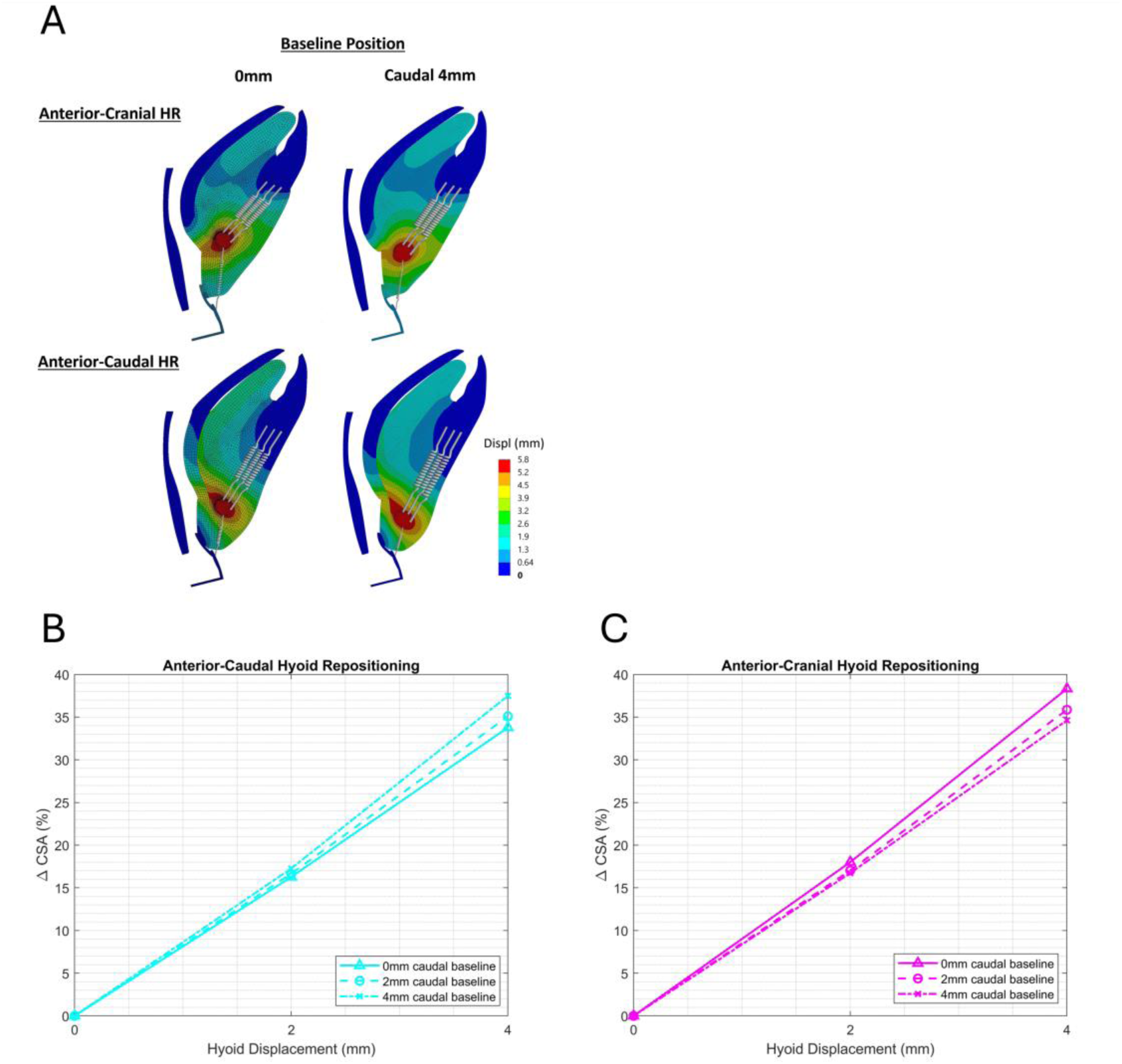
Tissue displacement and airway cross sectional area (CSA) for combined baseline hyoid position changes and surgical hyoid repositioning. (A) Color-coded resultant tissue displacement (Displ) fields shown at the maximal (4mm) surgical hyoid repositioning simulations in the anterior-cranial and anterior-caudal directions (similar to hyomandibular suspension and hyothyroidopexy surgeries, respectively) for a 4mm caudal hyoid baseline position (similar to the OSA phenotype). Percent change in CSA (ΔCSA) for surgical hyoid repositioning in (B) anterior-caudal and (C) anterior-cranial directions for 0, 2 and 4mm hyoid baseline position changes in the caudal direction. Both anterior-cranial and anterior-caudal surgical hyoid repositioning simulations produced large soft tissue displacement around the hyoid bone as well as the inferior portion of the soft palate and significantly increased the airway lumen CSA by ∼35%. Changes in the increment of hyoid baseline position produced a relatively small difference in ΔCSA. HR = hyoid repositioning.

#### Stress and strain

Figure 14 presents the stress and stain distributions across soft tissues with baseline hyoid position changes and surgical repositioning combined for the select directions and increments. The largest stresses and strains in upper airway soft tissues were located around the hyoid bone, in particular for anterior-cranial surgical hyoid repositioning simulations. The stress and strain distributions were almost identical for surgical hyoid repositioning from a 4mm caudal baseline hyoid position and the original baseline position.

**Figure 14:**
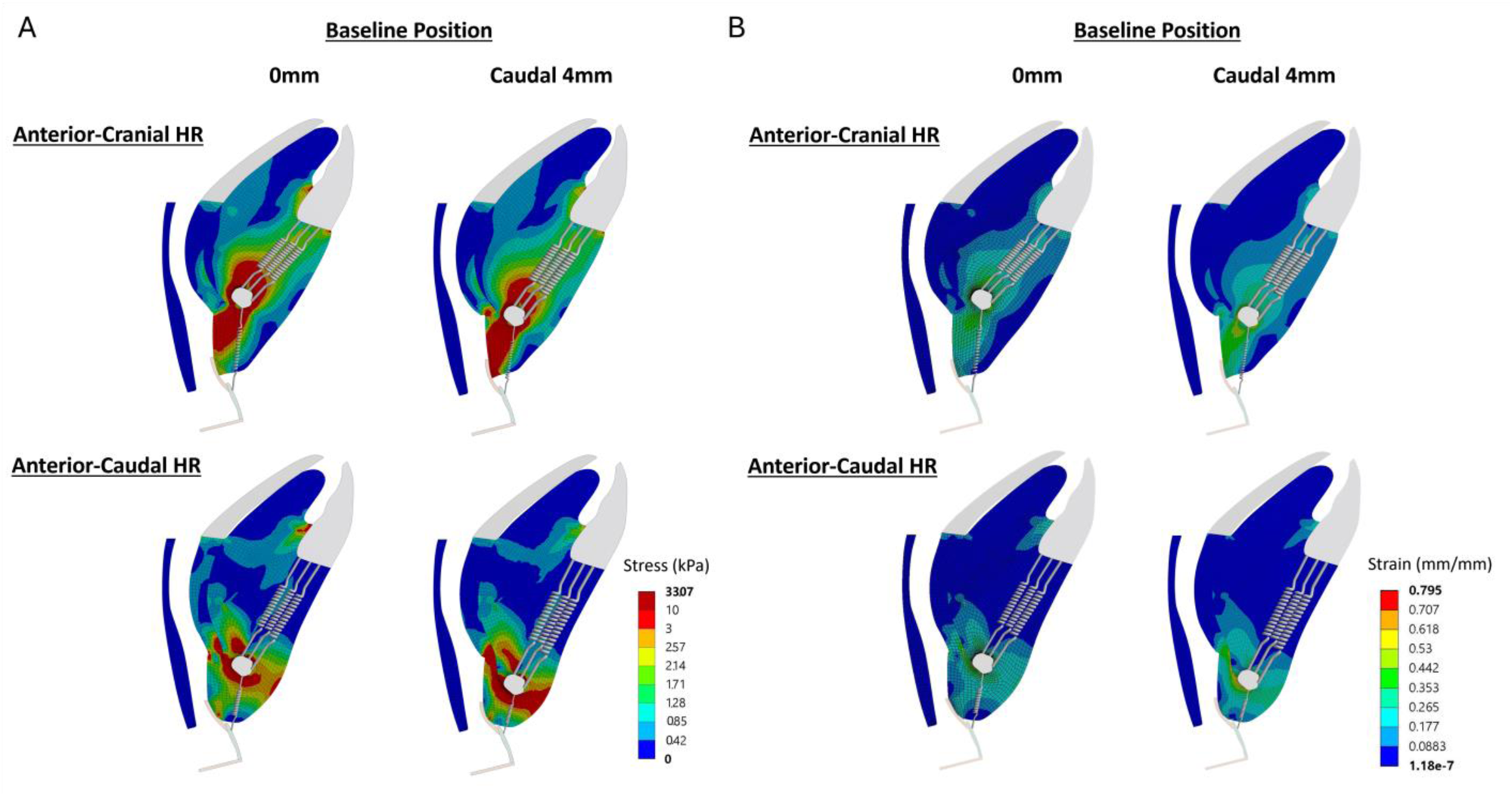
Tissue mechanics for combined hyoid baseline position changes and surgical hyoid repositioning. Stress (A) and strain (B) distribution results are shown for 4mm surgical hyoid repositioning simulations in the anterior-cranial and anterior-caudal directions for a 4mm caudal hyoid baseline position. Largest stresses and strains in upper airway soft tissues behind the hyoid bone were observed for anterior-cranial surgical hyoid repositioning. Note that only a subset of springs are shown. HR = hyoid repositioning.

## DISCUSSION

This is the first study to comprehensively investigate the role of the hyoid bone, both phenotype and surgical repositioning, on upper airway collapsibility, size and soft tissue mechanics. In summary, the model predicted that: 1) changes in the baseline hyoid position increase upper airway collapsibility regardless of the direction or increment of the new position; 2) anterior-based directions of surgical hyoid repositioning result in the largest improvement in upper airway collapsibility, lumen cross sectional area and tissue mechanics changes; and 3) the position of the hyoid at baseline influences the effectiveness of surgical hyoid repositioning in reducing upper airway collapsibility.

Particular strengths of this computational modeling study include: 1) it uniquely permitted the investigation of the effect on upper airway outcomes of baseline hyoid position, a phenotypic characteristic that cannot be directly studied in animal or human physiological models; 2) it was successfully validated against physiological outcomes to predict the percent change in upper airway closing pressure (Pclose) for surgical hyoid repositioning at various directions and increments; and 3) in addition to closing pressure, numerous outcomes such as upper airway lumen cross-sectional area, tissue displacement, stress and strain were computed, which allowed for a more comprehensive investigation into the role of the hyoid bone on upper airway patency and mechanics.

### Baseline hyoid position (phenotype) and upper airway collapsibility

The model predicted that changes in the baseline hyoid position led to an increase in upper airway collapsibility across all directions and increments. These findings suggest that the original baseline hyoid position represents the optimal upper airway anatomical configuration that minimizes upper airway collapsibility. This may be expected given that the original baseline position corresponds to the healthy rabbit airway anatomy. Changes to the hyoid bone’s baseline position may disrupt the balance of forces crucial for airway patency by increasing the area of unsupported tissues below, above or behind the hyoid bone, depending on the direction of the new position. Additionally, modifications in the length and orientation of muscles attached to the hyoid bone could affect passive muscle stretch during negative pressure simulations, diminishing their capacity to resist upper airway collapse and consequently increasing upper airway collapsibility (34).

Baseline hyoid position cannot be directly altered in animal or human physiological models. Nevertheless, several studies have investigated the association between baseline hyoid position, OSA and upper airway collapsibility in human participants. Those with OSA have been found to exhibit a significantly more inferior (or caudal) hyoid bone compared to healthy individuals (32, 35). Additionally, there exists a positive correlation between a more inferior hyoid position and increased severity of OSA, as indicated by the AHI (36). Furthermore, studies have shown a correlation between hyoid position and upper airway collapsibility, as assessed by the upper airway critical closing pressure (Pcrit) (37, 38). Specifically, a more caudal hyoid position is associated with a higher Pcrit, indicating increased upper airway collapsibility. These findings are consistent with the outcomes of the current study, where a 2-4mm caudal baseline hyoid position, akin to OSA, resulted in a ∼23-35% increase in Pclose (increased collapsibility). While factors such as age, BMI and neck circumference may potentially influence upper airway collapsibility in relation to hyoid baseline position in human studies, the current model uniquely demonstrates that anatomical changes specifically related to an altered hyoid baseline position impact upper airway collapsibility, irrespective of other factors.

### Surgical hyoid repositioning and upper airway outcomes

Our computational model suggests that the effect of surgical hyoid repositioning (from the original baseline hyoid position) on upper airway outcomes depends on the direction and increment of repositioning. Indeed, this is consistent with what we have previously reported in an anesthetized rabbit model for upper airway collapsibility (11).

### Anterior direction

The model predicted a large and progressive decrease in upper airway collapsibility with anterior surgical hyoid repositioning, with Pclose decreasing by 115% at 4mm. The model Pclose predications were in line with those achieved experimentally by Samaha et al (11). Model outcomes also support previous findings from other animal and human cadaver studies that have highlighted the beneficial impact of anterior hyoid repositioning on upper airway resistance and collapsibility (8–10).

In addition, our simulations demonstrated that anterior hyoid repositioning resulted in a large increase in the airway lumen cross sectional area accompanied by a high anterior displacement of the tongue, tissue mass and soft palate portion behind the hyoid bone. Displacement of upper airway soft tissues in response to surgical hyoid repositioning have not been quantified experimentally. However, select studies qualitatively observed the advancement of the tongue and widening of the velopharynx or hypopharynx in response to surgical hyoid advancement (8–10), supporting current model outcomes.

Since the soft tissues in the current model are defined as hyperelastic materials, an increase in stress or strain implies a stiffening of these tissues. Thus, anterior hyoid displacement led to a stiffening of soft tissues in a small area around the hyoid bone and in the lower portion of the soft palate. No previous studies have measured tissue stress or strain in response to anterior hyoid displacement (or any other direction for that matter), though stiffening of soft tissues around the hyoid bone may be expected as a result of stretch in the hyoid muscles. Compared to the other hyoid displacement directions, anterior hyoid displacement produced the least tissue stiffening.

### Cranial direction

Cranial hyoid repositioning led to a slight and progressive decrease in upper airway collapsibility in the current study. Apart from the experimental rabbit study by Samaha et al. (11), no studies have investigated the influence of cranial hyoid repositioning procedures on upper airway outcomes. Pclose predictions from the model were within the 95% confidence interval of the experimental results for all different magnitudes of cranial hyoid displacements. Nonetheless, the experimental study demonstrated no statistically significant impact of cranial repositioning on Pclose.

The model tissue displacement, stress and strain outcomes, as well as the upper airway cross sectional area, allow for a better understanding of the mechanisms underlying the Pclose outcome of cranial surgical hyoid repositioning. The first observation is that cranial hyoid displacement led to the cranial displacement of tissues below the hyoid bone and enlargement of the airway lumen in this region. Soft tissue stresses and strains are also increased below the hyoid, implying stiffening of upper airway walls. On the other hand, in the region above the hyoid bone, the soft tissues are compressed cranially, forcing the tongue and soft palate portion above the hyoid bone to fall back and narrow the airway lumen in that region. As a result, the overall lumen cross sectional area is unchanged after cranial hyoid repositioning. The slight improvement in Pclose may be attributed to stretching of infrahyoid tissues and muscles (e.g. thyrohyoid, sternohyoid springs), improving hyoid bone support during negative pressure application.

### Caudal direction

Caudal surgical hyoid repositioning resulted in a slight increase in upper airway collapsibility. While the model was largely in agreement with experimental findings, caudal hyoid repositioning did not yield any statistically significant impact on Pclose experimentally (11), the only other study to investigate this direction. Nonetheless, the experimental study revealed a general trend of a small increase in Pclose with caudal repositioning across many animals, indicating the possibility of this effect. Despite increases in stress/strain (and hence soft tissue stiffness) around the hyoid bone with caudal repositioning in the current model, the reduction in airway size, particularly evident below the hyoid, likely contributed to the airway collapsing at slightly lower negative pressures compared to baseline.

### Anterior-cranial and anterior-caudal directions

A relatively large and gradual decrease in upper airway collapsibility was predicted for anterior-cranial and anterior-caudal surgical hyoid repositioning by the model. These were at similar magnitudes to anterior repositioning, which is in agreement with the observations from the rabbit experiment study (11). Rosenbluth et al. (10) also detected an improvement in airway patency and stability in human cadavers with anterior-cranial and anterior-caudal hyoid displacement. No other studies have investigated the effect of hyoid displacement in these directions on upper airway collapsibility.

Anterior-cranial surgical hyoid repositioning produced the largest stress in tissues around the hyoid bone and specifically the airway wall in the region below the hyoid bone. These results suggest that anterior-cranial hyoid displacement stretches the tissues below the hyoid bone and infrahyoid muscles such as the sternohyoid and thyrohyoid leading to the enlargement of the airway lumen and the stiffening of the airway walls. Anterior-cranial surgical hyoid repositioning also led to the stiffening of a large portion of the tongue, mainly in the area between the hyoid and the mandible. Anterior-caudal hyoid displacement led to an increase in stress in the tissues at the boundary of the hyoid bone. Stresses of lower magnitudes were uniformly distributed in the soft palate tissues along the entire airway length indicating a moderate soft palate stiffening.

The major difference between anterior-cranial and anterior-caudal surgical hyoid displacements is in the resulting tissue stiffening produced. The stresses produced by anterior-caudal hyoid repositioning were more concentrated around the hyoid bone leading to the stiffening of a smaller region of soft tissues (mainly tissue mass located below the hyoid bone). On the other hand, anterior-cranial hyoid repositioning generated larger stresses that indicate higher stiffening of the soft tissues particularly in soft palate but also a wider spread of the stresses among the upper airway soft tissues. In fact, a large portion of the tongue tissue and the majority of the tissue mass is stiffened in response to anterior-cranial hyoid displacement. These observations lead to the proposition that anterior-cranial hyoid repositioning may result in further improvement in soft tissue mechanics than the other directions. Although the change in Pclose produced and the overall lumen cross-sectional area changes are similar, model predictions suggest that the manner in which anterior-cranial and anterior-caudal hyoid repositioning influence the upper airway differ.

### Baseline hyoid position effects on surgical hyoid repositioning and clinical implications

Model simulations suggest that upper airway collapsibility changes with surgical repositioning of the hyoid bone for the clinically relevant anterior-cranial and anterior-caudal directions (like hyomandibular suspension and hyothyroidopexy, respectively), depends on the initial baseline hyoid position. This highlights the importance of considering an individual’s baseline hyoid position when undertaking surgical repositioning procedures to achieve desired outcomes.

In cases where the hyoid was positioned more caudally, resembling the phenotype observed in OSA, the effectiveness of surgical hyoid repositioning in reducing upper airway collapsibility was reduced. Although the correlation between baseline hyoid position and surgical success of hyoid suspension surgeries has not been previously demonstrated, several studies have shown that the success rate of these surgeries is reduced with increased OSA severity (AHI), reflecting a more caudal baseline hyoid position (6, 33, 39). For instance, Vilaseca et al. (39) reported a surgical success rate for hyothyroidopexy procedure of 100% in their mild OSA group, 57% for moderate OSA and only 9% in the case of severe OSA. This may, at least in part, be related to the initial baseline hyoid position.

For the caudally positioned hyoid phenotype, the model predicts that 2mm anterior-cranial and anterior-caudal surgical hyoid repositioning led to a considerable decrease in Pclose by more than 50%. The model also showed that an overall increase in airway lumen cross-sectional area is produced by these simulations. These findings are consistent with the numerous studies that have demonstrated that hyoid suspension surgeries significantly decrease the AHI in most cases, although the surgical success rates are variable (5, 7). Note that surgical success of hyoid suspension procedures is generally defined as a 50% decrease in AHI and an AHI of less than 20 events/hr (6).

Both anterior-cranial and anterior-caudal surgical hyoid repositioning directions produced similar improvements in upper airway collapsibility and overall lumen enlargement with a more caudally located baseline hyoid position. However, there was a difference in the tissue displacements, stress and strain distributions induced by the two repositioning directions. Anterior-cranial surgical hyoid repositioning produced larger anterior displacements in the soft palate, particularly in the region behind the hyoid bone, and resulted in the advancement of epiglottis. The base of the tongue was also advanced with anterior-cranial hyoid displacement. A recent retrospective study by Van Tassel et al. (33) confirmed the increase in the airway size and anterior advancement of the tongue and epiglottis after hyomandibular suspension through endoscopic evaluation of the airway in awake and anesthetized patients. Other studies have also observed an advancement of the tongue and enlargement of the retro-epiglottic airspace in response to hyomandibular suspension (anterior-cranial direction) (6, 40). On the other hand, our model showed that anterior-caudal hyoid displacement pulls the tongue downwards and does not displace the epiglottis. In humans, a moderate increase in the airway size behind the base of the tongue has been observed after the hyothyroidopexy procedure, but the overall airway anatomical changes in response to the procedure were not significant (41). Moreover, anterior-cranial surgical hyoid repositioning led to larger soft tissue stiffening compared to anterior-caudal repositioning.

Model outcomes suggest that anterior-cranial surgical repositioning of the hyoid bone (e.g. hyomandibular suspension) may produce more favorable outcomes for OSA patients than anterior-caudal hyoid repositioning (e.g. hyothyroidopexy). Very few studies comparing hyomandibular suspension and hyothyroidopexy procedures have been undertaken. A small sample size study found that hyo-mandibular suspension produced larger reduction in the number of respiratory disturbances during sleep (apnea, hypopnea or respiratory event-related arousals) and higher improvement in the lowest oxygen saturation levels compared to hyothyroidopexy (42).

Apart from the more favorable improvement in airway geometry and tissue mechanics demonstrated in this study, hyomandibular suspension (with the improved technique introduced by Gillespie et al. (40)) presents several advantages compared to hyothyroidopexy procedure. In general, there is more room for hyoid displacement when suspending it to the mandible rather than the thyroid cartilage, i.e. larger displacement magnitude, specifically in the anterior direction.

Finally, results of the current study suggest that in addition to airway enlargement and tongue advancement, hyomandibular suspension is expected to stiffen the soft tissues in the hypopharynx region, increasing airway stability in this region. Ong et al. (6) showed that none of their OSA patients that had undergone hyomandibular suspension presented epiglottal collapse post-treatment. Consequently, the current study supports that hypopharyngeal and epiglottal collapse may be a potential useful patient selection criterion for hyomandibular suspension surgery.

Overall, our model findings underscore the importance of considering both the baseline hyoid position and the direction and extent of surgical repositioning in hyoid repositioning surgeries. Further research in humans will be required to ascertain the optimal hyoid position for repositioning, considering an individual’s natural baseline hyoid position. Nevertheless, our model outcomes lay the foundation for the pursuit of such investigations in human subjects.

### Limitations and Critique of Methods

Limitations related to the rabbit model and finite element model definitions, particularly boundary conditions, material properties and anatomical simplifications, have been detailed previously (17). Here we primarily limit discussion of limitations directly related to the current interventions.

### Closing pressure measurement

The absolute Pclose values obtained from the model are approximately 11 times larger than the physiological Pclose values (experimental baseline Pclose = −3.6 ± 0.9 cmH_2_O (11); model baseline Pclose = −40.15 cmH_2_O). The reason for this difference may be due to an over constrained upper airway, with boundary conditions applied to incorporate 3D behavior in 2D. Nonetheless, changes in outcomes with interventions was our primary concern. Indeed, changes in Pclose obtained from the model were found to successfully reproduce experimental changes in Pclose in response to surgical hyoid repositioning in five different directions and majority of the four magnitudes (11). To our knowledge, no other existing upper airway computational model validated their Pclose outcomes directly against experimental data in this manner (43–45).

While the model inherently has 3D functionality represented in 2D (17), the lateral upper airway walls are not represented, and upper airway collapse is solely in the antero-posterior direction. Thus, lateral wall collapse is not considered. Nonetheless, as discussed above, the model was still able to replicate the changes in Pclose found experimentally.

Lastly, the model encountered challenges in finding a Pclose solution for certain cases of combined baseline hyoid position and anterior-cranial/anterior-caudal surgical repositioning. Specifically, these cases included baseline hyoid positions in the cranial and/or anterior-cranial directions combined with 4mm anterior-cranial and anterior-caudal surgical repositioning (see Figure 12C,D). The substantial shifts in position, coupled with an intraluminal pressure load, caused excessive deformation that exceeded the model’s current capacity to accommodate. These magnitudes of repositioning and intraluminal pressure, in conjunction with the baseline hyoid shifts, represent extreme conditions for a rabbit and may potentially induce physiological tissue damage. Therefore, the model’s inability to converge on a solution may also reflect this.

### Baseline hyoid position (phenotype) changes

Human upper airway phenotypes with different hyoid baseline positions include multiple other differences in their airway anatomy related to facial skeletal patterns and tissue distribution (46–50). In the current model, different hyoid baseline position phenotypes have anatomical differences strictly related to the shift in hyoid bone location involving only structures that are in direct contact with the hyoid bone (inferior tongue edges, tissue mass and hyoid muscle connections). While it does not represent the different phenotypes assessed in clinical studies, the model is beneficial in examining the isolated effect of hyoid baseline position changes on upper airway collapsibility without other OSA confounding factors like the BMI, age, neck circumference, and fat pads in the neck region (32, 34, 51, 52).

### Future Directions of the Model

Future applications of this model should focus on exploring the impact of hyoid position—both in terms of phenotype and surgical repositioning—on the effectiveness of other interventions aimed at enhancing upper airway outcomes. These interventions could include crucial upper airway therapies or modifiers such as mandibular advancement, lung volume related tracheal displacement and upper airway dilator muscle activity. Outcomes from such simulations, including those from the current study, would provide proof-of-concept evidence for further investigation in humans.

## Conclusion

The upper airway computational finite element model in this study has allowed for the comprehensive investigation of the influence of baseline hyoid position (phenotype) and surgical hyoid repositioning on upper airway patency, collapsibility and tissue mechanics. The model successfully predicted changes in upper airway collapsibility in response to surgical hyoid repositioning interventions in multiple directions and increments. The model was also able to provide predictions of airway lumen cross-sectional area and pharyngeal tissue displacement, stress and strain distributions as a result of hyoid displacement that were qualitatively in agreement with literature. The original, healthy, baseline hyoid position was deemed the optimal upper airway anatomical configuration in terms of collapsibility. Furthermore, the position of the hyoid at baseline alters the effectiveness of surgical hyoid repositioning procedures in reducing upper airway collapsibility. Our model suggests that both baseline hyoid position and surgical hyoid repositioning direction and increment should be taken into consideration in surgical hyoid repositioning procedures for OSA. Study findings have potential implications to guide hyoid surgeries to improve OSA treatment outcomes and provide further insight into the hyoid’s role in OSA pathogenesis.

## APPENDIX A. Mesh Convergence

A mesh convergence study was performed to verify that the model is independent of the mesh density. The output used was the antero-posterior displacement of a portion of the soft palate located right behind the hyoid bone since it was found to be the site of collapse. Anterior-cranial surgical hyoid repositioning of 4mm magnitude was simulated for the mesh convergence study as it produced the largest tissue displacements and stresses and is a clinically relevant procedure (similar to hyomandibular suspension). Soft palate displacement outcomes were compared for multiple different mesh densities (see Figure A1 below). The mesh convergence study confirmed that for 3610 elements or higher, the model was independent of mesh density. Following mesh refinements surrounding the hyoid bone, the final mesh adopted consisted of 3970 elements (shown in Figure 1C). Table A1 summarizes the number of quadrilateral and triangular elements used for each mesh density used in Figure A1.

**Figure A1:**
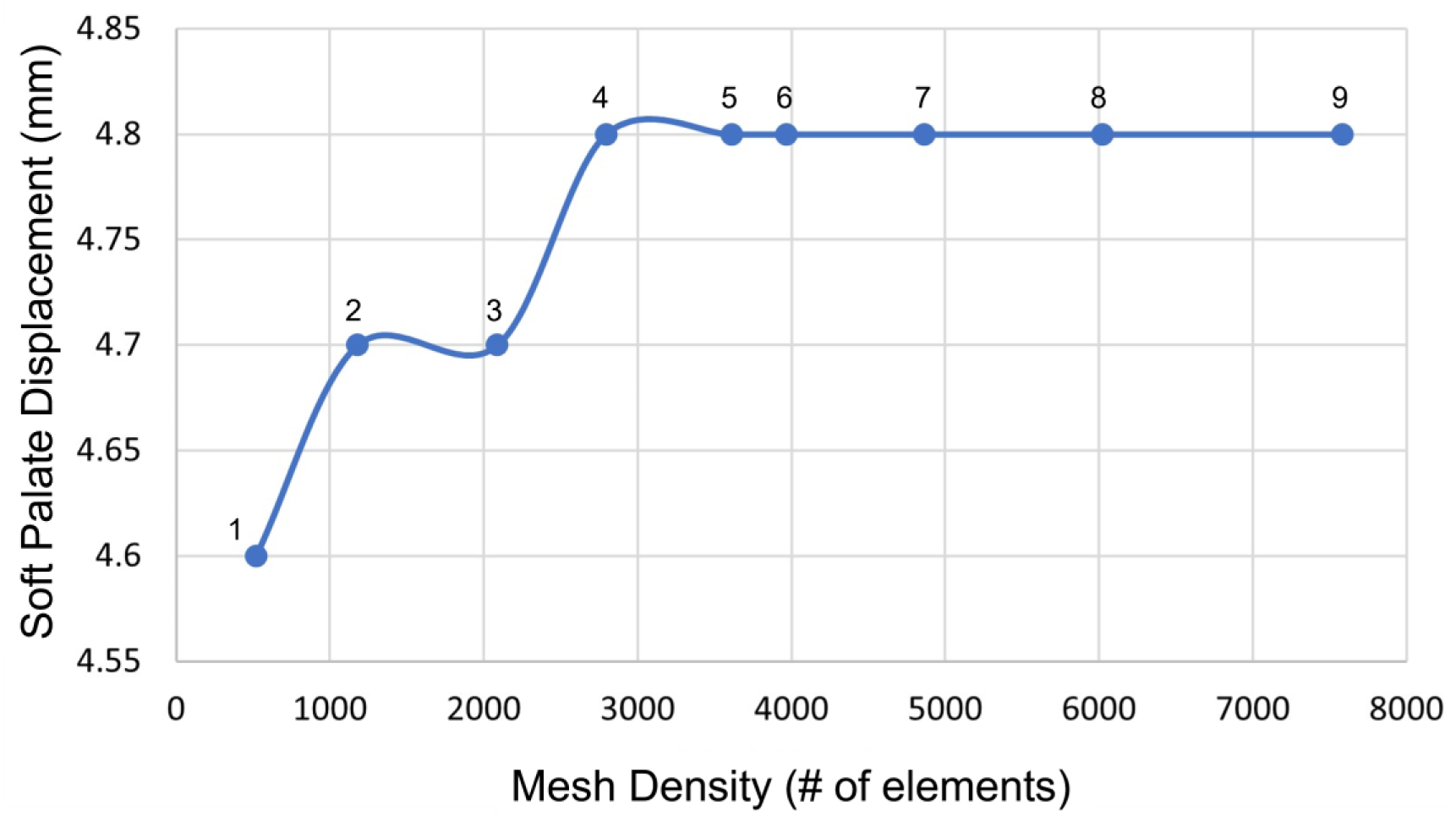
Mesh convergence study. Soft palate displacement in response to 4mm anterior-cranial surgical hyoid repositioning vs. mesh density (number of mesh elements). Convergence was reached at a mesh density of 3610 elements (point #5), with a final mesh consisting of refinements of 3970 elements used (see Figure 1C). Point numbers correspond to those in Table A1.

**Table A1:** Total mesh density, element types and numbers.

| **Point**<br>**#** | **Mesh Density**<br>**(Number of total**<br>**elements)** | **Number of**<br>**Quadrilateral**<br>**Elements** | **Number of**<br>**Triangular**<br>**Elements** |
| --- | --- | --- | --- |
| 1 | 519 | 487 | 32 |
| 2 | 1180 | 1123 | 57 |
| 3 | 2089 | 2031 | 58 |
| 4 | 2795 | 2739 | 56 |
| 5 | 3610 | 3553 | 57 |
| 6 | 3970<br>(final mesh) | 3907 | 63 |
| 7 | 4867 | 4811 | 56 |
| 8 | 6023 | 5944 | 79 |
| 9 | 7584 | 7498 | 86 |

Note that point number (#) corresponds to that in Figure A1.

## Acknowledgments

The authors would also like to thank Associate Professors Terence Amis and Kristina Kairaitis (The Westmead Institute for Medical Research and University of Sydney, Australia) and Professor Lynne Bilston (Neuroscience Research Australia and University of New South Wales, Australia) for their intellectual input and support.

## Data availability statement

Data supporting the results presented in this manuscript are shown in the manuscript figures and tables. Additional information can be made available upon to request to the authors.

## Author contributions

Model development and simulations were conducted at the American University of Beirut. JA conceived the research; DS and JA contributed to design of the research; DS advanced the model and performed the simulations; DS analyzed data; DS and JA interpreted the data; DS drafted the initial manuscript; DS and JA edited, revised, and approved final version of manuscript.

## REFERENCES

1. Peppard PE, Young T, Barnet JH, Palta M, Hagen EW, and Hla KM. Increased prevalence of sleep-disordered breathing in adults. American journal of epidemiology 177: 1006–1014, 2013.

2. Peppard PE, Young T, Palta M, and Skatrud J. Prospective study of the association between sleep-disordered breathing and hypertension. The New England journal of medicine 342: 1378–1384, 2000.

3. Young T, Finn L, Peppard PE, Szklo-Coxe M, Austin D, Nieto FJ, Stubbs R, and Hla KM. Sleep disordered breathing and mortality: Eighteen-year follow-up of the wisconsin sleep cohort. Sleep 31: 1071–1078, 2008.

4. Riley RW, Powell NB, and Guilleminault C. Obstructive sleep apnea and the hyoid: A revised surgical procedure. Otolaryngology--head and neck surgery : official journal of American Academy of Otolaryngology-Head and Neck Surgery 111: 717–721, 1994.

5. Song SA, Wei JM, Buttram J, Tolisano AM, Chang ET, Liu SY, Certal V, and Camacho M. Hyoid surgery alone for obstructive sleep apnea: A systematic review and meta-analysis. The Laryngoscope 126: 1702–1708, 2016.

6. Ong AA, Buttram J, Nguyen SA, Platter D, Abidin MR, and Gillespie MB. Hyoid myotomy and suspension without simultaneous palate or tongue base surgery for obstructive sleep apnea. World J Otorhinolaryngol Head Neck Surg 3: 110–114, 2017.

7. Kezirian EJ, and Goldberg AN. Hypopharyngeal surgery in obstructive sleep apnea: An evidence-based medicine review. Archives of Otolaryngology–Head & Neck Surgery 132: 206–213, 2006.

8. Graaff WBVd, Gottfried SB, Mitra J, Lunteren Ev, Cherniack NS, and Strohl KP. Respiratory function of hyoid muscles and hyoid arch. 57: 197–204, 1984.

9. Benderro GF, Gamble J, Schiefer MA, Baskin JZ, Hernandez Y, and Strohl KP. Hypoglossal nerve stimulation in a pre-clinical anesthetized rabbit model relevant to osa. Respir Physiol Neurobiol 250: 31–38, 2018.

10. Rosenbluth KH, Kwiat DA, Harrison MR, and Kezirian EJ. Hyoid bone advancement for improving airway patency:Cadaver study of a magnet-based system. 146: 491–496, 2012.

11. Samaha CJ, Tannous HJ, Salman D, Ghafari JG, and Amatoury J. Role of surgical hyoid bone repositioning in modifying upper airway collapsibility. Frontiers in Physiology 13: 2022.

12. Yang Z. Finite element analysis for biomedical engineering applications. CRC Press, 2019.

13. Srodon PD, Miquel ME, and Birch MJ. Finite element analysis animated simulation of velopharyngeal closure. Cleft Palate Craniofac J 49: 44–50, 2012.

14. Huang Y, White DP, and Malhotra A. The impact of anatomic manipulations on pharyngeal collapse: Results from a computational model of the normal human upper airway. Chest 128: 1324–1330, 2005.

15. Xu C, Brennick MJ, Dougherty L, and Wootton DM. Modeling upper airway collapse by a finite element model with regional tissue properties. Med Eng Phys 31: 1343–1348, 2009.

16. Ilegbusi OJ, Kuruppumullage DNS, Schiefer M, and Strohl KP. A computational model of upper airway respiratory function with muscular coupling. Computer Methods in Biomechanics and Biomedical Engineering 25: 675–687, 2022.

17. Amatoury J, Cheng S, Kairaitis K, Wheatley JR, Amis TC, and Bilston LE. Development and validation of a computational finite element model of the rabbit upper airway: Simulations of mandibular advancement and tracheal displacement. Journal of applied physiology (Bethesda, Md : 1985) 120: 743–757, 2016.

18. Amatoury J, Kairaitis K, Wheatley JR, Bilston LE, and Amis TC. Peripharyngeal tissue deformation and stress distributions in response to caudal tracheal displacement: Pivotal influence of the hyoid bone? Journal of applied physiology (Bethesda, Md : 1985) 116: 746–756, 2014.

19. Amatoury J, Kairaitis K, Wheatley JR, Bilston LE, and Amis TC. Peripharyngeal tissue deformation, stress distributions, and hyoid bone movement in response to mandibular advancement. 118: 282–291, 2015.

20. Wingerd BD. Rabbit dissection manual. Baltimore: Johns Hopkins University Press, 1985.

21. Kirkness JP, Christenson HK, Garlick SR, Parikh R, Kairaitis K, Wheatley JR, and Amis TC. Decreased surface tension of upper airway mucosal lining liquid increases upper airway patency in anaesthetised rabbits. J Physiol 547: 603–611, 2003.

22. Amatoury J, Kairaitis K, Wheatley JR, Bilston LE, and Amis TC. Peripharyngeal tissue deformation and stress distributions in response to caudal tracheal displacement: Pivotal influence of the hyoid bone? J Appl Physiol 116: 746–756, 2014.

23. Brouillette RT, and Thach BT. Control of genioglossus muscle inspiratory activity. J Appl Physiol 49: 801–808, 1980.

24. Amatoury J, Kairaitis K, Wheatley JR, Bilston LE, and Amis TC. Peripharyngeal tissue deformation, stress distributions, and hyoid bone movement in response to mandibular advancement. J Appl Physiol 118: 282–291, 2015.

25. Amatoury J, Cheng S, Kairaitis K, Wheatley JR, Amis TC, and Bilston LE. Development and validation of a computational finite element model of the rabbit upper airway: Simulations of mandibular advancement and tracheal displacement. J Appl Physiol 120: 743–757, 2016.

26. Kairaitis K, Stavrinou R, Parikh R, Wheatley JR, and Amis TC. Mandibular advancement decreases pressures in the tissues surrounding the upper airway in rabbits. Journal of Applied Physiology 100: 349–356, 2006.

27. Kairaitis K, Verma M, Amatoury J, Wheatley JR, White DP, and Amis TC. A threshold lung volume for optimal mechanical effects on upper airway airflow dynamics: Studies in an anesthetized rabbit model. Journal of Applied Physiology 112: 1197–1205, 2012.

28. Olson L, Ulmer L, and Saunders N. Pressure-volume properties of the upper airway of rabbits. Journal of Applied Physiology 66: 759–763, 1989.

29. Kirkness JP, Eastwood PR, Szollosi I, Platt PR, Wheatley JR, Amis TC, and Hillman DR. Effect of surface tension of mucosal lining liquid on upper airway mechanics in anesthetized humans. J Appl Physiol 95: 357–363, 2003.

30. Schiefer M, Gamble J, Baskin J, and Strohl K. Hypoglossal nerve stimulation in a rabbit model of obstructive sleep apnea reduces apneas and improves oxygenation. J Appl Physiol 129: 442–448, 2020.

31. Serghani M-M, Heiser C, Schwartz AR, and Amatoury J. Exploring hypoglossal nerve stimulation therapy for obstructive sleep apnea: A comprehensive review of clinical and physiological upper airway outcomes. Sleep Med Rev 76: 101947, 2024.

32. Chi L, Comyn FL, Mitra N, Reilly MP, Wan F, Maislin G, Chmiewski L, Thorne-FitzGerald MD, Victor UN, Pack AI, and Schwab RJ. Identification of craniofacial risk factors for obstructive sleep apnoea using three-dimensional mri. The European respiratory journal 38: 348–358, 2011.

33. Van Tassel J, Chio E, Silverman D, Nord RS, Platter D, and Abidin MR. Hyoid suspension with uppp for the treatment of obstructive sleep apnea. Ear, nose, & throat journal 1455613211001132–1455613211001132, 2021.

34. Bilston LE, and Gandevia SC. Biomechanical properties of the human upper airway and their effect on its behavior during breathing and in obstructive sleep apnea. Journal of applied physiology (Bethesda, Md : 1985) 116: 314–324, 2014.

35. Barrera JE, Pau CY, Forest VI, Holbrook AB, and Popelka GR. Anatomic measures of upper airway structures in obstructive sleep apnea. World J Otorhinolaryngol Head Neck Surg 3: 85–91, 2017.

36. Bilici S, Yigit O, Celebi OO, Yasak AG, and Yardimci AH. Relations between hyoid-related cephalometric measurements and severity of obstructive sleep apnea. J Craniofac Surg 29: 1276–1281, 2018.

37. Genta PR, Schorr F, Eckert DJ, Gebrim E, Kayamori F, Moriya HT, Malhotra A, and Lorenzi-Filho G. Upper airway collapsibility is associated with obesity and hyoid position. Sleep 37: 1673–1678, 2014.

38. Sforza E, Bacon W, Weiss T, Thibault A, Petiau C, and Krieger J. Upper airway collapsibility and cephalometric variables in patients with obstructive sleep apnea. Am J Respir Crit Care Med 161: 347–352, 2000.

39. Vilaseca I, Morelló A, Montserrat JM, Santamaría J, and Iranzo A. Usefulness of uvulopalatopharyngoplasty with genioglossus and hyoid advancement in the treatment of obstructive sleep apnea. Archives of Otolaryngology–Head & Neck Surgery 128: 435–440, 2002.

40. Gillespie MB, Ayers CM, Nguyen SA, and Abidin MR. Outcomes of hyoid myotomy and suspension using a mandibular screw suspension system. Otolaryngology--head and neck surgery 144: 225–229, 2011.

41. Stuck BA, Neff W, Hörmann K, Verse T, Bran G, Baisch A, Düber C, and Maurer JT. Anatomic changes after hyoid suspension for obstructive sleep apnea: An mri study. Otolaryngology--head and neck surgery 133: 397–402, 2005.

42. Mickelson SAMDFA. Hyoid advancement to the mandible (hyo-mandibular advancement). Operative techniques in otolaryngology--head and neck surgery 23: 56–59, 2012.

43. Huang Y, Malhotra A, and White DP. Computational simulation of human upper airway collapse using a pressure-/state-dependent model of genioglossal muscle contraction under laminar flow conditions. Journal of applied physiology (Bethesda, Md : 1985) 99: 1138–1148, 2005.

44. Huang Y, White DP, and Malhotra A. The impact of anatomic manipulations on pharyngeal collapse: Results from a computational model of the normal human upper airway. Chest 128: 1324–1330, 2005.

45. Liu Y, Mitchell J, Chen Y, Yim W, Chu W, and Wang RC. Study of the upper airway of obstructive sleep apnea patient using fluid structure interaction. Respir Physiol Neurobiol 249: 54–61, 2018.

46. Gündüz Arslan S, Dildeş N, and Devecioglu Kama J. Cephalometric investigation of first cervical vertebrae morphology and hyoid position in young adults with different sagittal skeletal patterns. TheScientificWorld 2014: 159784–159788, 2014.

47. Haralabakis NB, Toutountzakis NM, and Yiagtzis SC. The hyoid bone position in adult individuals with open bite and normal occlusion. European journal of orthodontics 15: 265–271, 1993.

48. Jena AK, and Duggal R. Hyoid bone position in subjects with different vertical jaw dysplasias. The Angle orthodontist 81: 81–85, 2011.

49. Mortazavi S, Asghari-Moghaddam H, Dehghani M, Aboutorabzade M, Yaloodbardan B, Tohidi E, and Hoseini-Zarch S-H. Hyoid bone position in different facial skeletal patterns. Journal of clinical and experimental dentistry 10: e346–e351, 2018.

50. Sheng C-M, Lin L-H, Su Y, and Tsai H-H. Developmental changes in pharyngeal airway depth and hyoid bone position from childhood to young adulthood. The Angle orthodontist 79: 484–490, 2009.

51. Young T, Skatrud J, and Peppard PE. Risk factors for obstructive sleep apnea in adults. Jama 291: 2013–2016, 2004.

52. Bayat M, Shariati M, Rakhshan V, Abbasi M, Fateh A, Sobouti F, and Davoudmanesh Z. Cephalometric risk factors of obstructive sleep apnea. CRANIO® 35: 321–326, 2017.

